# Macrophage depletion restores the DRG microenvironment and prevents axon degeneration in bortezomib-induced neuropathy

**DOI:** 10.1101/2025.01.22.634362

**Authors:** Michael B. Thomsen, Abhishek Singh, Christina N. Thebeau, Vivian D. Gao, Nicholas F. Schulze, Oshri Avraham, Sarah X. Yang, Shriya Koneru, Sami S. Geier, Shannon M. Landon, Aidan Pelea, Valeria Cavalli, Stefanie Geisler

**Author notes:** These authors contributed equally to this work (Abhishek Singh, Christina N. Thebeau). Department of Cellular Biology, University of Georgia; Athens, USA.

## Abstract

Peripheral neuropathy is a common and debilitating side effect of the chemotherapeutic bortezomib (BTZ). To explore the mechanisms underlying BTZ-induced neuropathy (BIPN), we developed a mouse model that replicates the route of administration and approximates the prolonged BTZ exposure experienced by patients. We find that male mice treated with BTZ experience more severe sensorimotor dysfunction and axon loss compared to females and observed similar results when analyzing human data. Using single cell RNA-sequencing, we reveal that BTZ significantly alters the dorsal root ganglia (DRG) microenvironment in mice, producing pronounced sex-specific changes in satellite glial cells (SGCs) in males and females and dysregulation of the extracellular matrix (ECM), particularly in males. These changes are accompanied by expansion of macrophages, which is more pronounced in males. We identify four macrophage subtypes in the DRG, including a pro-fibrotic population that is exclusively associated with BIPN. Depletion of macrophages via anti-CSF1R treatment in male mice prevents BTZ-induced SGC activation and aberrant collagen deposition in DRGs, potently preserves peripheral axons, and improves functional outcomes. These findings highlight SGCs, neuroinflammation and dysregulation of the ECM as drivers of sex-specific differences in BIPN and suggest that targeting neuroinflammation is a promising therapeutic strategy to treat this disease.

**One Sentence Summary:** Inflammation and neurofibrosis in DRGs underlie sex-specific differences in bortezomib-induced neuropathy and are promising treatment targets.

## INTRODUCTION

The introduction of bortezomib (BTZ) for the treatment of multiple myeloma and other hematologic cancers in 2003 transformed the therapeutic landscape, leading to significantly higher complete remission rates and improved overall survival outcomes (*1*). To this day, BTZ is the drug most used to treat multiple myeloma (*2*). Unfortunately, the benefits of BTZ are limited by side effects. About half of all patients treated with BTZ develop BTZ-induced neuropathy (BIPN), and up to a third of patients require dose reductions that diminish therapeutic efficacy and overall survival (*3–8*). BIPN is characterized by sensorimotor dysfunction, including pain, tingling, numbness, and distal weakness, which can lead to ambulatory difficulties sufficient to require use of a wheelchair (*6, 9, 10*). These profoundly disabling symptoms can be associated with anxiety, depression, and loss of productivity – all leading to decreased quality of life - and may constrain continuation of treatment (*11, 12*). Current strategies to ameliorate BIPN are limited to treating positive symptoms, such as pain and tingling symptomatically, while negative symptoms, such as numbness and imbalance and the underlying mechanisms of BIPN cannot be targeted due to a lack of mechanistic insights. In contrast to most BTZ-induced side effects, BIPN can last long after treatment has ended.

BTZ inhibits the 20S core proteasome, which results in cancer cell death via multiple mechanisms, including suppression of the unfolded protein response, induction of reactive oxygen species and preventing the degradation of various pro-apoptotic factors (*13*). BTZ does not penetrate the central nervous system but crosses the blood-nerve barrier to accumulate in dorsal root ganglia (DRG) and peripheral nerves, which consequently are the main anatomical sites of direct neurotoxicity (*14, 15*). While axon degeneration in some chemotherapy-induced neuropathies occurs axon-autonomously, we and others have shown that BTZ-induced degeneration is transcriptionally regulated and thus requires signals from the cell body (*16, 17*). However, what these signals are and how they arise is unknown.

Cell bodies of sensory peripheral nerves are located within the DRG, where they are fully enveloped by satellite glial cells (SGCs). This unique arrangement allows SGCs to serve as essential regulators of neuronal activity, homeostasis, and repair (*18*). Other cells within the DRG microenvironment include endothelial cells, fibroblasts, immune cells and Schwann cells, which, as actively dividing cells, are also susceptible to the effects of BTZ, perhaps more so than neurons.

However, the extent to which the DRG microenvironment is involved in axonal injury in peripheral neuropathies generally, and in BIPN specifically, is largely unknown. To address this gap in knowledge we developed a mouse model of chronic BIPN and used single cell RNA sequencing and immunohistochemistry to interrogate the DRG microenvironment. We find that BIPN causes more severe neuropathy in males than females and confirmed this finding in human patients. Axonal pathophysiology in the mouse model is associated with profound changes in the DRG microenvironment. The greatest transcriptional response occurred in SGCs in male mice. Upregulated genes encoding component of the extracellular matrix (ECM) were associated with massively increased collagen deposition and macrophage expansion in DRGs, more in males than females. Depleting macrophages in males normalized the DRG microenvironment, potently prevented axon degeneration, and improved functional outcomes. Accordingly, targeting the DRG microenvironment may be an effective strategy to treat BIPN.

## RESULTS

### BIPN in mice is more severe in males

BTZ is mostly administered intravenously in patients, frequently for several months up to two years and sometimes beyond (*2*). Patients commonly start to experience neuropathy symptoms around the 4^th^ cycle, typically about 3.5 months out (*5, 9, 19*). To approximate this long-time course, we treated a cohort of female and a cohort of male mice (n = 10 mice per group) with intravenous injections of 0.8 mg/kg BTZ or vehicle twice weekly for eight weeks (Fig. 1A) and confirmed the initial findings in two additional, separate female and male cohorts (n=10-15 mice per group; Fig. 1A). Data were analyzed for each cohort separately and then combined. If different results were obtained between the cohorts, the data are reported for each cohort.

**Figure 1:**
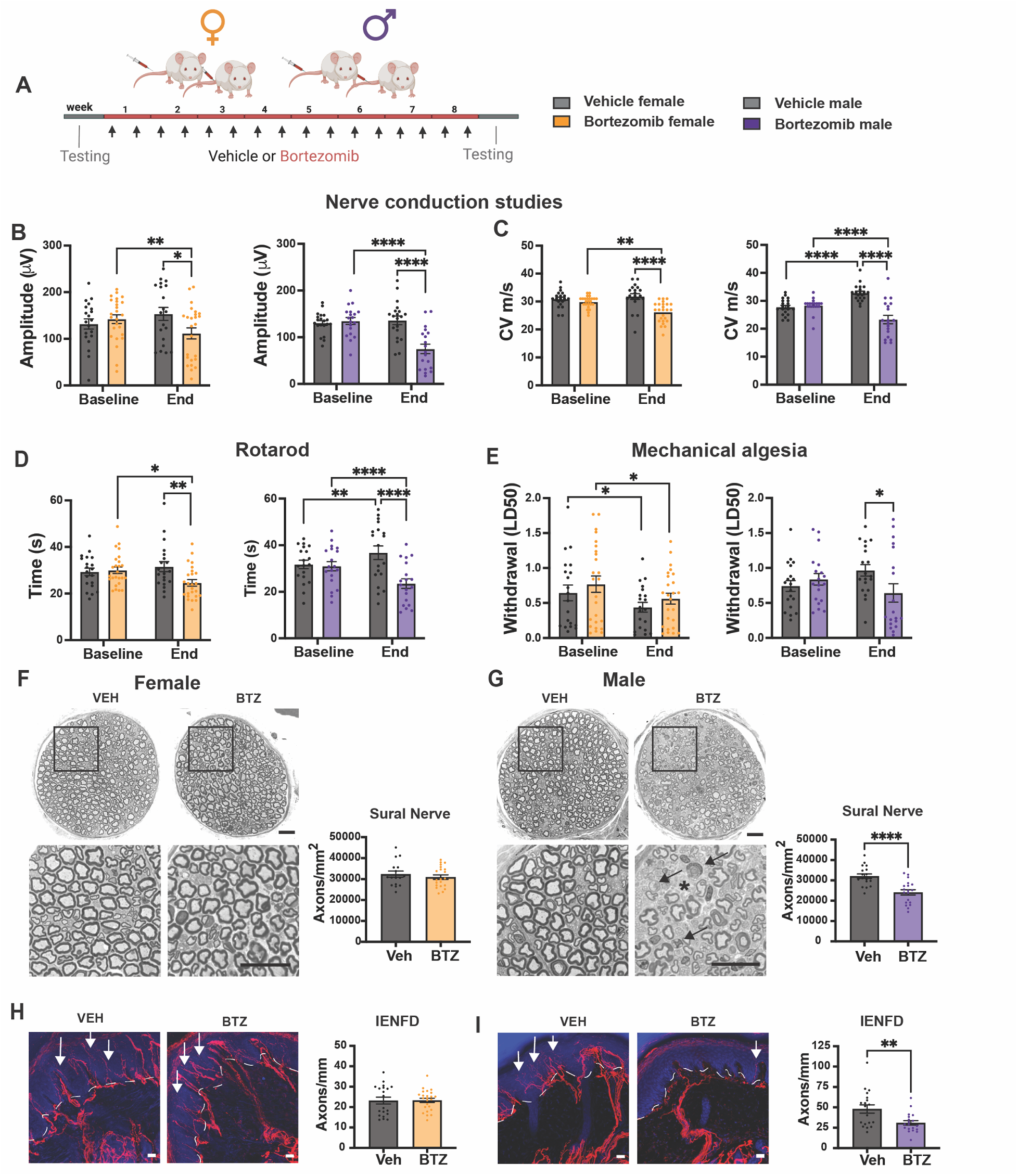
BIPN is more severe in male than female mice. (**A)** Experimental layout. (**B, C**) Nerve conduction studies of the tail nerve. The tail compound nerve amplitude potentials (B) and conduction velocities (C) decrease significantly in animals treated with BTZ as compared to baseline measurements and vehicle-treated mice. Two-way ANOVA, followed by Fisher’s LSD B - female: *P=0.0124; **P=0.0029; male ****P<0.0001; C – female **P=0.0012; ****P<0.0001; male - ****P<0.0001; n=19-25 mice per group. (**D**) Female and male mice treated with BTZ stayed on the Rotarod for a shorter time period as compared to baseline measurements and vehicle-treated mice. Two-way ANOVA, followed by Fisher’s LSD; female *P=0.0103, **P=0.0036; male **P=0.0055, ****P<0.0001; n=19-25 mice per group. (**E)** When compared to vehicle-treated mice, the mechanical withdrawal threshold is significantly reduced after BTZ treatment in male, but not female mice. Two-way ANOVA, followed by Fisher’s LSD female-Veh baseline vs end *P=0.0282, BTZ baseline vs end *P=0.0142; male *P=0.0195; n=19-25 mice per group. (**F, G)** Photomicrographs of toluidine-blue stained semithin cross sections of the distal sural nerve at low (upper row) and high (lower row) magnifications from female (F) and male (G) mice treated with vehicle or BTZ. (F) There is no difference in axon morphology and density between vehicle and BTZ-treated female mice. Unpaired t-test P=0.9446. (G) Axons appear less densely packed (asterisk), and some axons express signs of axon degeneration (arrows), resulting in decreased axon density in BTZ-treated male mice. Unpaired t-test, ****P<0.0001. Scale bars 20 μm. (**H, I)**: Nerve fibers stained with the pan-axonal marker PGP 9.5 (red; arrows) extend into the epidermis (nuclei stained with DAPI in blue). There is no difference in intra-epidermal nerve fiber density (IENFD; arrows) between vehicle- and BTZ-treated female mice (H), unpaired t-test P=0.9466. In contrast, IENFD is reduced in male mice treated with BTZ as compared to vehicle treated mice (I). Unpaired t-test **P=0.0052; Scale bars 20 μm.

Patients with BIPN have more involvement of large, myelinated axons than small, thinly myelinated and unmyelinated nerve fibers (*5*). We therefore evaluated large fiber dysfunction by performing nerve conduction studies in the tail, which revealed a significant decrease in the compound nerve action potential (CNAP) amplitude in mice treated with BTZ as compared to baseline and to the vehicle group with a greater decline in males than females (Fig. 1B). Conduction velocity was also reduced in both sexes, suggesting loss of the largest and fastest conducting fibers, an effect that, again, was more pronounced in males than females (Fig. 1C).

BTZ-induced large fiber dysfunction in humans can result in numbness and imbalance. To evaluate for sensory-motor dysfunction, mice were tested on the rotarod. Male and female mice treated with BTZ were unable to stay on the rotarod as long as before BTZ treatment was initiated and as long as vehicle treated groups (Fig. 1D). BIPN in patients is frequently painful and can be associated with mechanical hyperalgesia (*20*). Using a graded series of von Frey filaments, we detected a significantly lowered mechanical withdrawal threshold in male mice as compared to vehicle-treated mice (Fig 1E). In contrast, the mechanical withdrawal threshold in vehicle and BTZ-treated female mice was not different (Fig. 1E), suggesting that mechanical hyperalgesia had develops in male, but not female, BTZ-treated mice.

We next assessed whether the observed functional deficit observed in BTZ-treated mice is associated with distal axon degeneration and axon loss. Because BIPN in patients is sensory predominant and symptoms start distally in the feet (*17*), we analyzed myelinated axons of the sural nerve, a distal sensory nerve that is frequently used for nerve biopsies in human patients. We found signs of axon degeneration and axon loss in male, but not female, mice treated with BTZ (Fig. 1F, G). The modest decrease in CNAP amplitude in female mice without frank axon loss may indicate damaged, but not (yet) lost large axons or that dysfunction at the level of the soma contributes to the observed large fiber dysfunction in females. Pain and temperature signals are conveyed by unmyelinated C-fibers and thinly myelinated A-δ fibers, which are routinely evaluated clinically in skin biopsies taken from the ankle by staining fixed sections with the pan-axonal marker PGP9.5 to assess intraepidermal nerve fiber density (IENFD) (*21*). We found that male mice treated with BTZ had a significantly reduced IENFD as compared to those treated with vehicle (Fig. 1I). In contrast, there was no difference in the IENFD between vehicle and BTZ treated female mice (Fig. 1H). Thus, our mouse model of chronic BIPN shows sensory predominant deficits affecting large and small nerve fibers that were more pronounced in male than female mice.

### BIPN is more severe in men than women

To the best of our knowledge, sex differences of BIPN have not been described previously in human patients. To evaluate whether the severity of BTZ-induced neuropathy differs between male and female patients we analyzed data of patients diagnosed with BIPN by a Neuromuscular specialist from the Peripheral Neuropathy Research Registry (*22*) and from our own tertiary care center neuropathy clinic. Male (n=24) and female (n=16) patients with BIPN had similar age and co-morbidities (Table S1). Sensory nerve conduction studies of the sural nerve revealed that significantly more men than women with BIPN had absent or reduced sural sensory nerve action potentials (SNAPs), consistent with more myelinated axon impairment and/or loss in men (Figure 2A). Dysfunction of myelinated axons in patients can be examined by testing sensation to vibration. In accord with the greater reduction in sural SNAP amplitude, more men than women with BIPN had absent or reduced sensation to vibration at the toes and ankles (Fig. 2B, C). Sensation to pin at the toes and ankles, which tests C and A-δ fibers, was similar in men and women with BIPN (Fig. 2D, E), whereas sensation to pin at the fingers was reduced in more men than women (Fig. 2F). Neuropathic pain intensity was similar in men and women suffering BIPN, with 65% of men and 63% of women reporting intense or moderate pain (Fig. 2G). Pain intensities in other peripheral neuropathies and pain conditions are often higher in females (*23–26*), which suggests relatively increased pain intensity in males with BIPN. Finally, we re-analyzed data of a previously published cohort that contained both the sex and total neuropathy score (TNSc) of each individual patient treated with BTZ (*9*). We found that men had a significantly higher TNSc than women, indicating more severe neuropathy in men (Fig. 2H). The current data validate a greater severity of BIPN in male than female patients. Replication of the same findings in mice strengthens the clinical relevance of the animal model.

**Figure 2:**
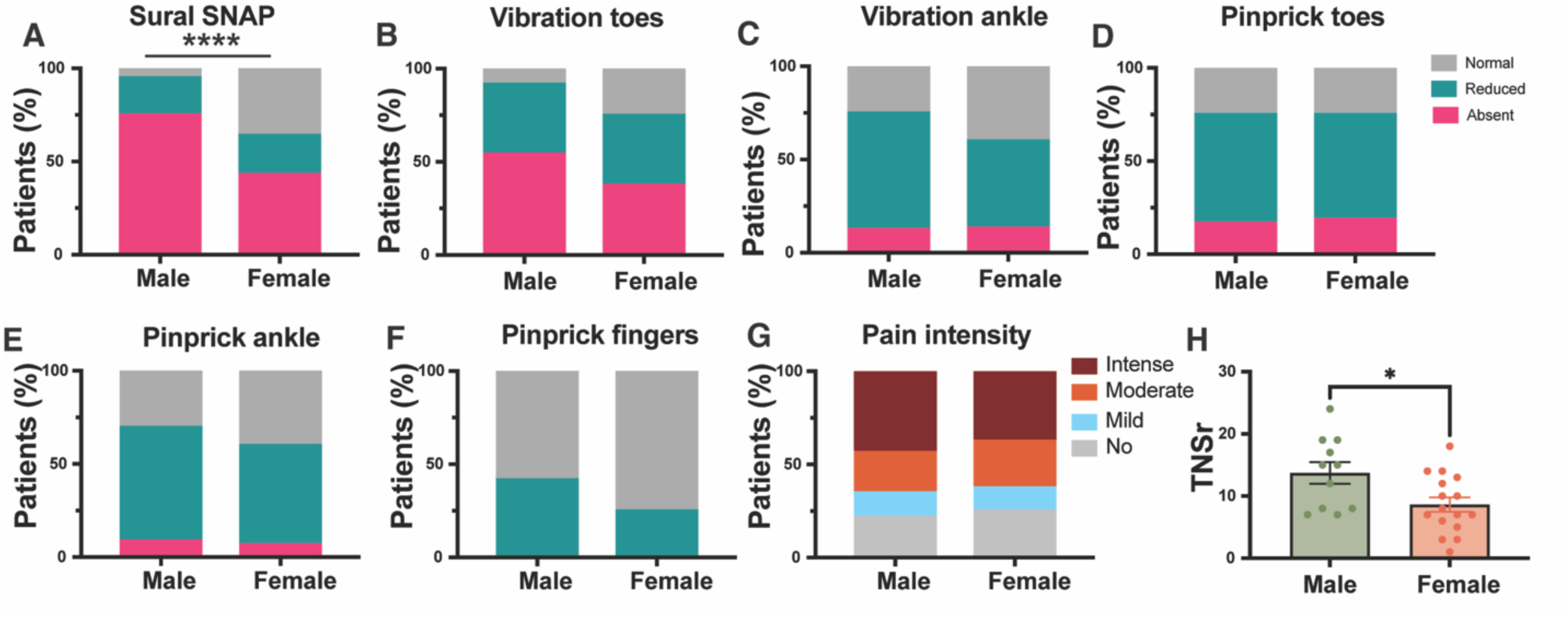
BIPN is more severe in men than women. (**A-F)**: Percent of patients diagnosed with BIPN exhibiting absent (magenta), reduced (cyan) or normal (gray) sensory nerve action potential (SNAP) of the sural nerve (A), sensation to vibration at the toes (B) and ankles (C), and sensation to pinprick at the toes (D), ankles (E), and fingers (F). Chi square test **** P<0.0001; male n=24, female n=16. (**G)**: Percent of patients with BIPN experiencing intense (pain intensity 7-10 on a scale from 0-10), moderate (intensity 4-6 out of 10), mild (intensity 1-3 out of 10) or no pain; male n=24, female n=16. (**H)**: Analysis from Chaudry et al. (*9*) showing increased total neuropathy score (TNSr) in male patients with BIPN. Data ± standard error of the mean; unpaired t-test *P=0.0182; male n=11, female n=16.

### ScRNA seq reveals expansion of non-neuronal cells in DRGs of mice with BIPN

The finding of fiber dysfunction without axon loss in females suggests that changes at the level of the neuronal soma contribute to the phenotype. Furthermore, preclinical studies by us and others have demonstrated that BTZ-induced axon degeneration, at least in part, is transcriptionally regulated in the neuronal cell body (*16, 17*). The somata of peripheral sensory nerves reside in DRG, which also contain several kinds of non-neuronal cells. Because non-neuronal cells are replicating cells, they are potentially more vulnerable to the action of a proteasome inhibitor, such as BTZ, and may contribute to the observed phenotype. To gain insights into the impact of BTZ on the DRG microenvironment, we carried out single cell RNA sequencing of lumbar DRGs from female and male mice treated with BTZ or vehicle (Figs. 1A, 3A). We obtained more than 10,000 cells per condition for a total of 62,063 high-quality single cell transcriptomes (Supplemental file 1). Using reciprocal PCA to integrate samples by batch, we identified ten non-neuronal and seven neuronal transcriptionally distinct cell clusters in the DRG that were characterized by robust markers of gene expression and consistent with previous DRG cell type annotations (*27–29*) (Fig. 3B, D, S1; Supplementary data file 2). Among the cells identified, SGCs, marked by expression of *Ptprz1*, *Fabp7*, and *Kcnj10* (Fig. 3D, S1D), constituted the largest cell population in the DRG (38.9 % of all cells), followed by non-myelinating Schwann Cells (16.3 %) (Fig. 3B). Neurons, distinguished by expression of canonical markers including *Tubb3*, *Isl1*, and *Snap25* (Fig. S1C) constituted about 10.2% of DRG cells, and included nociceptors I (*Trpv1*) and II (*Calca, Ntrk1)*, mechanoreceptors (*Ntrk2*), cold neurons (*Trpm8*), itch neurons (*P2rx3*), C light touch mechanoreceptors C-LTMR (*Th*), and proprioceptors (*Pvalb;* Fig. 3B, D, S1C).

**Figure 3:**
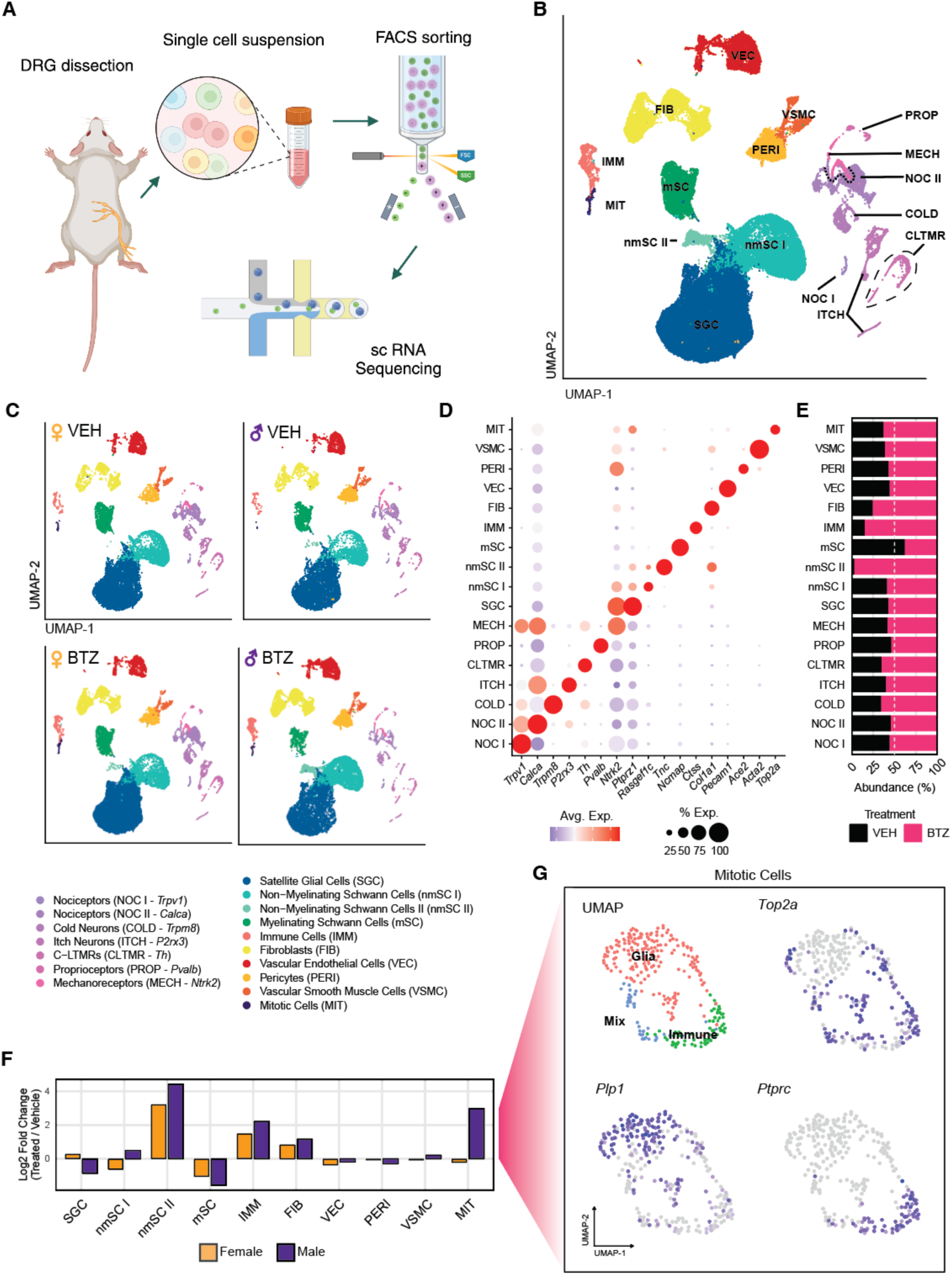
Single cell RNA sequencing of DRGs identifies neuronal and non-neuronal cells. **(A)** Schematic of experimental design. (**B)** UMAP plot of 62,063 DRG cells from male and female treated with BTZ or vehicle (n=4 per sex and condition) identifies seven neuronal and 10 non-neuronal cell types. (**C)** UMAP plots of all cells separated by sample condition (**D)** Dot plot of selected marker genes specific to each major cell type cluster. (**E)** Relative abundance of DRG cell types in BTZ-treated vs. veh-treated samples. (**F)** Normalized cell abundance changes in non-neuronal cell populations following BTZ treatment in male and female samples showing a relative increase in non-myelinating Schwann cells, immune cells and mitototic cells of DRGs from females (yellow) and males (purple) following BTZ. (**G)** UMAP of independently clustered mitotic cells (Top2A) reveals three groups, which express cell markers of glial (PlP1) and immune (Ptprc) markers.

To determine whether the cell composition of DRGs changed following BTZ treatment, we compared the identified clusters between vehicle and BTZ conditions (Fig. 3C). Separate UMAP plots for sex and treatment showed good congruence, revealing the same 17 cell clusters in vehicle and BTZ treated animals, but with a shift of relative cell counts in some clusters following BTZ administration (Fig. 3C, E, F). The myelinating Schwann cell was the only cell type that was relatively decreased by BTZ, especially in male mice (Fig. 3E, F), in accord with the observed loss of myelinated axons following BTZ in our model. In contrast, mitotic cells, comprising clusters with gene expression signatures of glial and immune lineages (Fig. 3G), as well as mature immune cells, non-myelinating Schwann cells, and fibroblasts showed the largest relative increases in abundance (Fig. 3C, E, F). These data suggest a shift of the cellular composition of the DRG in BIPN, characterized by an expansion of glial and immune cells and an overall loss of mature myelinating Schwann cells.

### SGCs undergo profound changes in response to BTZ in both sexes, with a greater magnitude in males

To gain insights into how BTZ alters DRG cell types at the transcriptional level, we overlaid the numbers of differentially expressed genes (DEGs) onto the UMAP plots for each condition subset (Fig. 4A). We focused our initial analysis on neuronal subpopulations (Fig. S2) and noticed varied responses to BTZ. Surprisingly, several neuron types in DRGs of both sexes expressed few if any DEGs following chronic BTZ, including mechanoreceptors (*Ntrk2*), cold neurons (*Trpm8*), and nociceptors 1 (*Trpv1;* Fig. 2SA). The greatest gene expression changes following BTZ treatment were observed in P2rx3-positive “itch” neurons and nociceptors II (*Calca*) in both sexes, whereas increased gene expression in C-light touch mechanoreceptors (C-LTMR, *Th*) and proprioceptors (*Parv)* were seen in females, but not males (Fig. 2SA). Gene ontology analysis revealed especially strong upregulation of genes involved in lipid metabolism in C-LTMRs from females with BIPN (Fig. S2B), whereas the same genes were downregulated in several neuron types of males (Fig S2C). In contrast, genes encoding components of the extracellular matrix (ECM) were increased in several neuron types in males (Fig. S2E) but largely unchanged in females (Fig S2D), suggesting that neuron populations from males and females engage distinct signaling pathways in response to BTZ. Overall, however, neuronal subtypes expressed relatively few DEGs following BTZ, when compared to non-neuronal cells (Fig. 4A).

**Figure 4:**
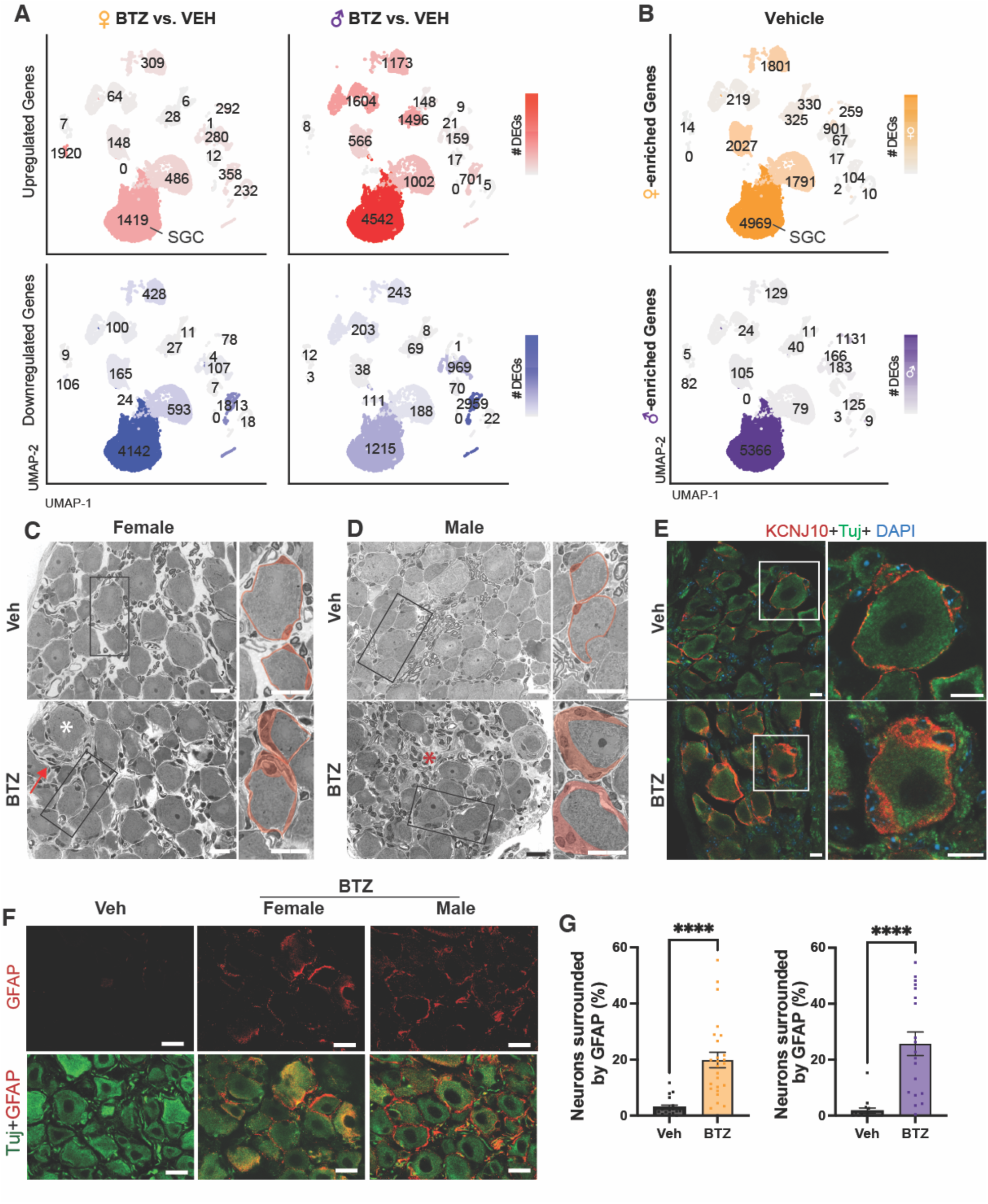
Satellite glia cells are reactive in BIPN. (**A)** UMAP plots displaying differential gene expression magnitude between vehicle and BTZ treated female and male mice in all major cell type clusters (p-adj < 0.05). Red: upregulated, Blue: downregulated. (**B)** UMAP plots displaying differential gene expression magnitude between female and male vehicle samples. p-adj < 0.05. (**C, D**) Photomicrographs of toluidine blue stained semithin sections of lumbar DRG from female (C) and male (D) mice treated with vehicle or BTZ. Boxed area enlarged to the right of the photos. SGCs pseudo-colored in red. A rare neuron (white asterisk) is surrounded by increased interstitium (red arrow) in female mice treated with BTZ (C), whereas the interstitium (red asterisk) is massively increased in males with BIPN (D). Scale bars 20 μm (**E)** Photomicrographs of DRGs from males with BIPN stained with anti-tubulin (green) to mark neurons and KCNJ10 (red) to label SGCs reveal widening of the perineuronal sheet after BTZ. Scale bars 10 μm. (**F**) Photomicrographs of DRGs stained with TUJ (green, neurons) and GFAP (red, reactive satellite glial cells) from vehicle- and BTZ-treated female and male mice. Note the increase of GFAP staining after BTZ. Scale bars 20 μm. (**G**) Quantification of neurons surrounded by GFAP-ir in female and male mice treated with veh or BTZ. Unpaired t-test ****P<0.0001 (n=16-25 mice per group).

The greatest increase of DEGs genes following BTZ was observed in SGCs (Fig. 4A). Although SGCs expressed the most DEGs in both males and females with BIPN, the pattern of gene expression differed between both sexes (Fig. 4A). In male SGCs, more genes were up- than down-regulated in response to BTZ, whereas the opposite was true for SGCs from females (Fig. 4A; p<0.0001; Fisher’s t-test). We therefore wondered whether the SGCs baseline transcriptome differs between males and females. Indeed, of all cell types in the DRG, SGCs had the largest set of sex-specific DEGs (Fig. 4B). In addition to enrichment of X chromosome-specific transcripts, female SGCs showed enrichment of genes involved in lipid metabolism (*Idi1*, *Mvd*, *Lss)* and transport (*Mfsd2a* and *Slc27a1;* Fig. S3A). In contrast, male SGCs had higher expression of genes related to protein processing and cellular catabolic processes, including *Uchl1*, *Hspb1*, and *Cd24a* (Fig. S3A). Accordingly, gene ontology analysis revealed enrichment of pathways involved in proteasome function and assembly in male SGCs (Fig. S3B). This could render SGCs in males more susceptible to the degenerative actions of the proteasome inhibitor BTZ.

SGCs completely envelop DRG neurons thereby providing metabolic and trophic support, modulating neuronal activity and playing a key role in sensory transmission (*18*). Given the importance of the neuron-glia interactions, we investigated whether the BIPN-induced transcriptional response in SGCs resulted in morphological alterations. We observed a thin perineuronal layer of SGCs around each neuron in semithin sections of DRGs stained with toluidine-blue from vehicle treated mice (Fig. 4C, D), as has been described before (*30*). Following treatment of female mice with BTZ, the appearance of many SGCs was indistinguishable from those treated with vehicle, apart from a few neurons that were surrounded by a widened perineuronal layer (Fig. 4C). In male BTZ-treated mice, however, we frequently observed prominent widening of the perineuronal layer, suggesting profound SGC hypertrophy (Fig. 4D). Staining of DRG sections from BTZ-treated male mice with Fabp7 and Kcnj10, which are established markers for SGCs (*31*) revealed widening of immune-positive rings surrounding neurons, confirming marked hypertrophy of BTZ-treated SGCs in males (Figs. 4E, S3C-E).

In response to nerve injury and in different pain conditions, the expression of Glial fibrillary acidic protein (GFAP) is increased in SGCs (*18*). Here, we find increased expression of *Gfap* mRNA, more prominent in males than females, in the SGCs of mice with BIPN (Fig. S3F). We confirmed increased protein expression by counting the number of neurons surrounded by GFAP-immunoreactivity (Fig. 4F). While in vehicle treated mice, about 2% of neurons were surrounded by GFAP-immunoreactive cells, the number of neurons enveloped by GFAP-positive cells was increased more than ten times in both male and female BTZ-treated mice (Fig. 4 F, G), indicating increased reactivity of SGCs in male and female mice with BIPN.

Thus, SGCs exhibit pronounced transcriptional changes that are greater in males than females with BIPN. Additionally, male mice with BIPN exhibit significant SGCs hypertrophy, a characteristic not observed in female mice.

### SGCs in males, but not females, increase expression of extracellular matrix components in response to BTZ

In line with increased SGC reactivity in BIPN as demonstrated by increased GFAP-expression and SGC hypertrophy, expression of several immediate-early gene transcription factors, including *Atf3*, *Fos* and *Jun*, was increased in SGCs of mice with BIPN (Fig. 5A, B). The expression was increased in both females and males but was significantly greater in males (Fig. 5C). In addition, the expression of several genes engaged in cell cycle regulation and migration were elevated in both males and females, including cell cycle checkpoint protein cyclin D2 (Ccnd2) and the receptor tyrosine kinase ephrin A5 (Epha5; Fig. 5A-C).

**Figure 5:**
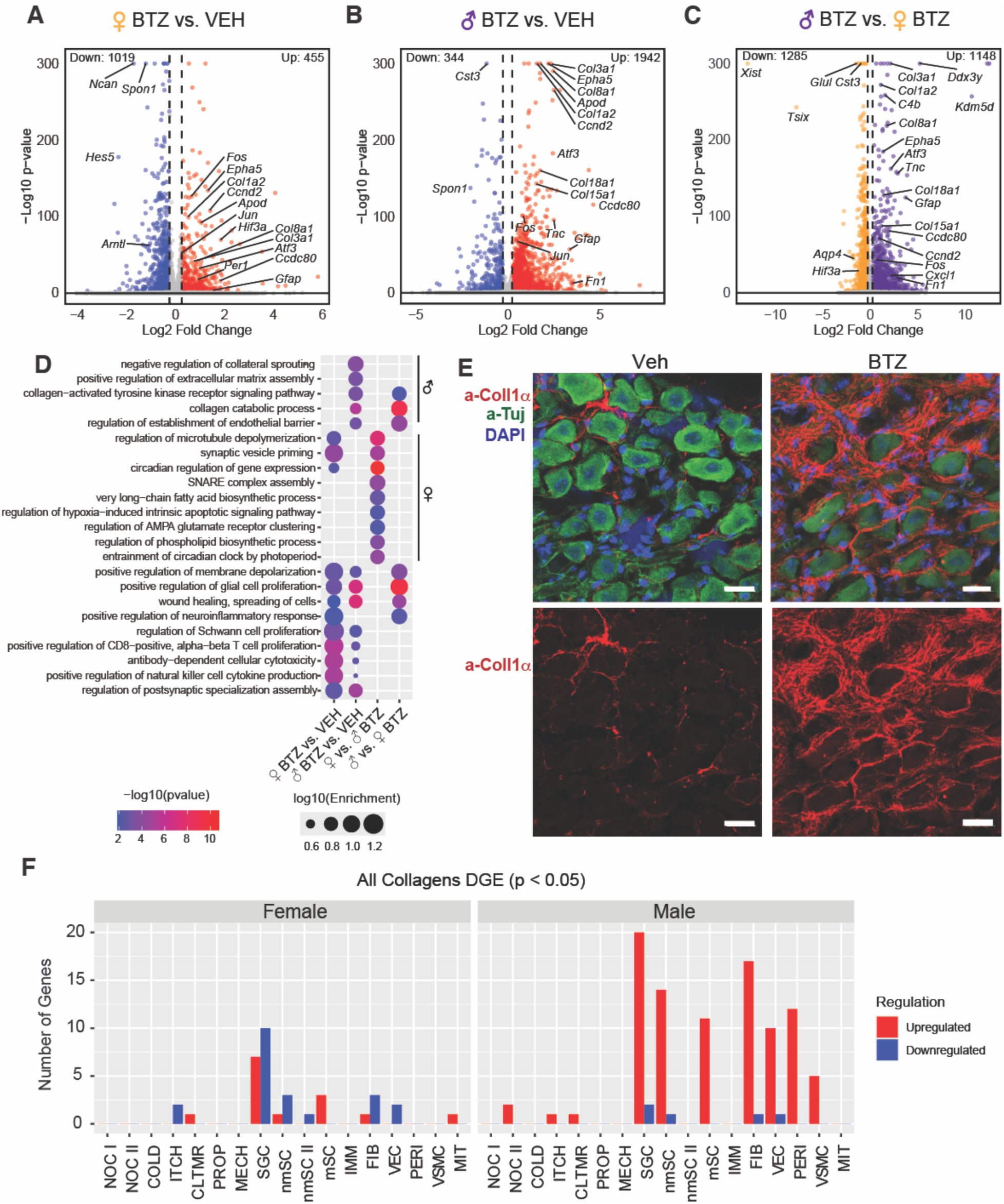
Neurofibrosis in DRGs of male mice with BIPN. (**A, B**) Volcano plots of differential gene expression in SGCs of female (A) and male (B) vehicle and BTZ-treated mice (red: upregulated, p-ad j< 0.05, log2FC > 0.25; blue: downregulated, p-adj < 0.05, log2FC <-0.25) show increased expression of immediate early genes, and some ECM genes with greater magnitude in males. (**C**) Volcano plot of differential gene expression in SGCs between male and female BTZ-treated samples (purple male enriched: p-adj< 0.05, log2FC > 0.25; yellow: female enriched: p-adj < 0.05, log2FC <-0.25) showing significantly higher expression of immediate early genes and ECM in males, whereas Hif3alpha is increased in female SGC with BIPN. (**D**) Gene Ontology enrichment analysis of differentially expressed genes (p-adj < 0.05, log2FC > 0.25) in SGCs from male and female mice treated with vehicle or BTZ. **(E)** Photomicrographs of DRGs stained for tubulin (TUJ, green, neurons), collagen 1 alpha (Coll1α, red) and DAPI (nuclear marker, blue) from males treated with vehicle (left) or BTZ (right). Scale bars: 20 μm. **(F)** Number of significantly up-(red) or down-(blue) regulated collagen genes (out of 46) per cell type in females (left) and males (right) with BIPN.

Surprisingly, among the most upregulated genes in SGCs of mice with BIPN were those encoding proteins of the extracellular matrix (ECM). Expression of some of these genes were upregulated in both sexes with greater increases in males, while others were upregulated exclusively in males (Fig. 5A-C). Collagen I (*Col1a2)*, III *(Col3a1)* and VIII (*Col8a1)* and the ECM component *Ccdc80* (coiled-coil domain containing 80), which plays an important role in ECM organization, were increased in both sexes (Fig. 5A, B), but significantly more enriched in males than females treated with BTZ (Fig. 5C). Collagens XV (*Col15)* and XVIII (*Col18)*, Tenascin-C (*Tnc*), a component of perineuronal nets comprising a specialized form of ECM, and fibronectin (*Fn1*) a glycoprotein that binds to membrane-spanning integrins, were strongly upregulated in male SGCs but not changed in females with BIPN (Fig. 5B, C). Accordingly, GO analysis (biological function) revealed enrichment of pathways associated with ECM organization, including collagen catabolic processes, wound healing and collagen-activated TRK signaling pathways only in SGCs of male mice with BIPN (Fig. 5D).

While SGCs of males with BIPN were characterized by prominent changes in genes encoding ECM proteins, SGCs of females with BIPN showed differential expression of core circadian clock components such as *Per1* and *Arntl* (Fig. 5A, C). GO analysis (biological function) revealed enrichment of pathways relating to "circadian rhythms", in SGC of females with BIPN, which were not present in male SGCs (Fig. 5D). Female DEGs also were uniquely enriched for terms related to lipid metabolism and neurotransmission (Fig. 5D). These findings indicate that BTZ elicits distinct responses in male and female SGCs, with males upregulating ECM-related genes and females modulating expression of genes associated with circadian regulation, lipid metabolism, and neurotransmission.

### Neurofibrosis develops in DRGs of males with BIPN

Because male mice expressed a more severe phenotype, we explored the ECM induced changes of male mice in more detail at the protein level. Collagen Iα immunoreactivity around the SGCs-neuron units was greatly increased in male mice treated with BTZ compared to vehicle (Fig 5E). As pathological increase and remodeling of ECM in peripheral organs is referred to as fibrosis and a similar process in the brain was recently termed neurofibrosis (*32*), we refer to the excessive collagen deposition in DRGs of mice with BIPN as neurofibrosis. Considering the profound increase of collagen Iα in DRGs of mice with BIPN, we wondered whether cells other than SGCs in DRGs also increase expression of genes encoding ECM components in response to BTZ. Expression analysis of 46 different collagen genes across all cell types showed that SGCs have the greatest number of upregulated collagen genes in males treated with BTZ (Fig. 5F). In BTZ-treated male mice, other cell types with increased collagen expression include fibroblasts, non-myelinating Schwann cells, pericytes, myelinating Schwann cells and vascular endothelial and smooth muscle cells (Fig. 5F). Remarkably, many cell types in female DRGs treated with BTZ respond with a prominent downregulation of genes encoding different collagens (Fig. 5F), extending the sex-specific differences in ECM expression from SGCs to the other cell types in the DRG.

Components of the ECM do not only have structural functions but are also involved in inter-cellular communication (*33–35*). To evaluate if collagen contributes to inter-cellular communication in the DRG, we turned to cell chat analysis in R, a computational tool used to infer and analyze cell-cell communication between groups of cells using single cell transcriptomics data and ligand-receptor interaction information (*36*). We found that collagen was the top signaling molecule in SGCs and fibroblasts (Fig S4A, B), suggesting that BTZ-induced dysregulation of the ECM will disrupt intercellular communication in DRGs.

### Immune cell populations expand in DRGs of mice with BIPN, more in males than females

Increased ECM is frequently the result of persistent immune activation (*37, 38*). Our initial analysis revealed expansion of immune cell populations in DRGs of mice with BIPN (Fig. 3C, F). To gain insights into which immune cells change in response to BTZ, we performed unbiased clustering and gene analysis of the immune cell populations separately. We identified four distinct macrophage populations, dendritic cells, T-cells, B-cells and neutrophils in the DRGs (Fig 6A). We also observed a cluster of cells that expressed marker genes of both macrophages (*Maf, C1qa, C1qb*) and SGCs (*Fabp7, Ptprz1*; Fig 6A, C). These cells were recently identified as Imoonglia (*39*) and have been shown to also increase in numbers after peripheral and central nervous system injuries (*40*).

**Figure 6:**
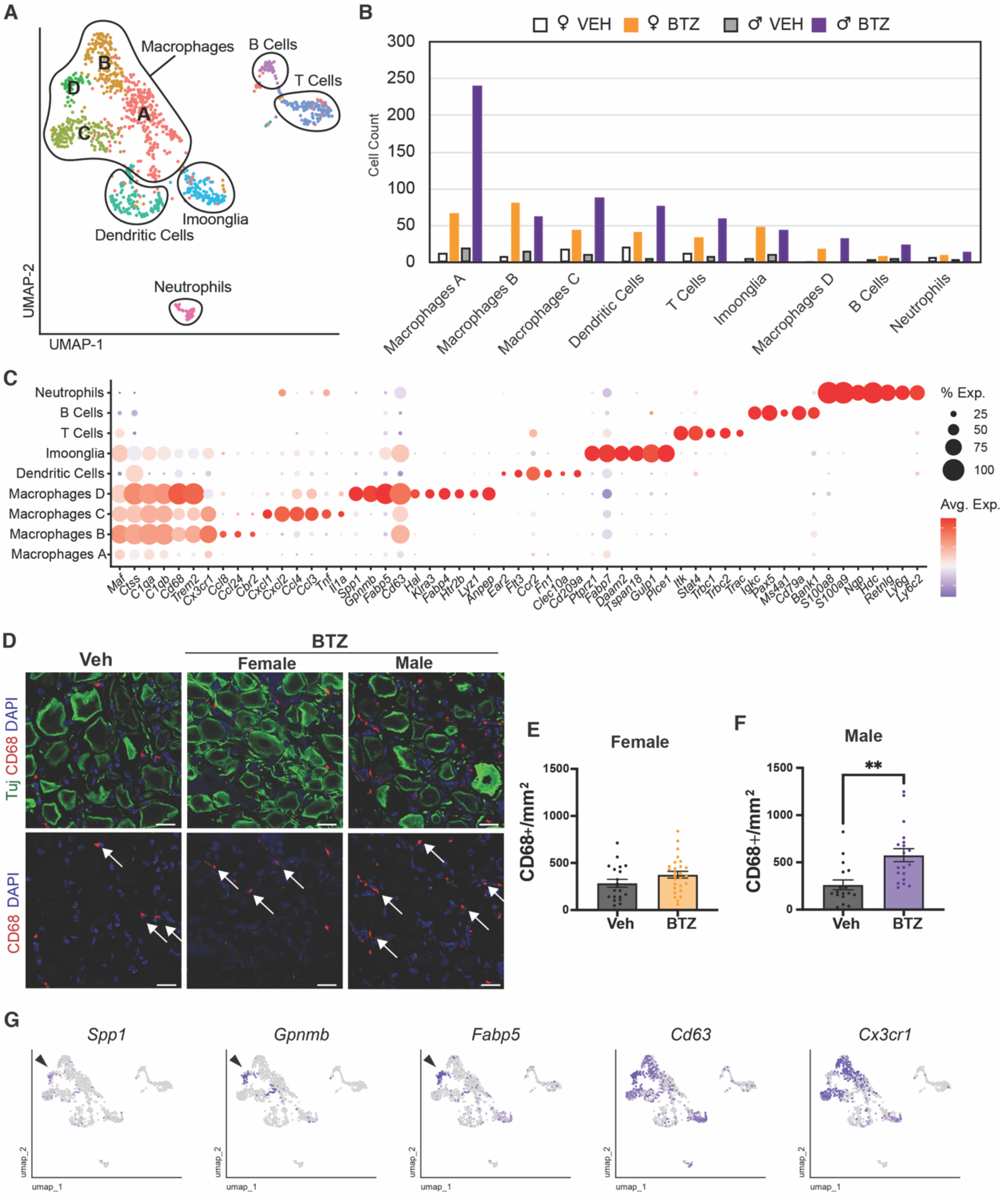
Immune cell population abundance is increased in DRGs following BTZ. (A) UMAP plot identifies 6 immune cell populations in the mouse DRG. (**B**) Absolute abundance of immune cell subpopulations in DRGs of vehicle and BTZ-treated female and male mice. (**C)** Dot plot of selected enriched marker genes for each immune cell subpopulation. **(D)** Photomicrographs of DRGs stained for tubulin (Tuj, green, neurons), CD68 (red, macrophages) and DAPI (blue, nuclei) from female and male mice treated with vehicle or BTZ. Arrows point to CD68-positive cells. Scale bars: 20 μm. **(E, F)** Quantification of CD68-positive cells in DRGs of females (E) and males (F) treated with vehicle or BTZ. Unpaired t-test **P=0.0012; n = 17-25 mice per group. (**G**) UMAP plots of marker genes of scar-associated macrophages in immune cell populations. Arrows point to macrophage-D cluster.**F**

Following BTZ treatment, numbers of many immune cell types were elevated in DRGs, generally more in males than females (Fig. 6B). Among these, macrophages showed the largest increase (Fig. 6B). To confirm macrophage expansion in DRGs, we stained sections with the common macrophage marker CD68 (Fig. 6D), which is strongly expressed by three of the four macrophage types (Fig. 6C). In females, CD68 positive macrophages significantly increased in one cohort, but not the other, resulting overall in a non-significant trend of increased CD68 positive cells in DRGs of females with BIPN (Fig. 6 D, E). In males, numbers of macrophages increased significantly in both cohorts following BTZ (Fig. 6D, F), consistent with the greater expansion of macrophages in BTZ-treated males as seen in our transcriptomic data (Fig. 6B).

Analysis of the macrophage cluster revealed four distinct cell populations. Macrophage C expressed strong pro-inflammatory markers, including *Il1a, Tnf, Cxcl1, Ccl3* and *Ccl4* (Fig 6C). Macrophage D is *Trem2*-positive and expressed genes that have been associated with scar-associated macrophages, including *Spp1*, *Gpnmb*, *Fabp5*, and *Cd63* (*41*) (Fig. 6C, G). Scar associated macrophages have been recently shown to be broadly fibrogenic in different diseases in humans, including liver cirrhosis and lung fibrosis (*41*). Interestingly, this macrophage type is not present in the DRG of vehicle treated mice (Fig. 6B), but emerges in BIPN, with greater increase in males than in females. CellChat analysis revealed macrophages, particularly macrophage subtype D, as the cell type in DRGs that exhibits the most pronounced differential interaction strengths in BIPN (Fig. S4C, D). Macrophage D received increased signals from SGCs and fibroblasts and showed enhanced interactions with most cell types, especially with other macrophages, SGCs and immonglia in BIPN (Fig S4D).

Thus, the male-predominant DRG immune response in BIPN is characterized by marked expansion of pro-inflammatory and scar-associated macrophage populations, which significantly increased their interactions with other cell types, including SGCs.

### Depletion of macrophages normalizes DRG microenvironment and protects from axon degeneration in males

Fibrosis in peripheral organs and the CNS is the result of chronic inflammation (*37, 38*). Blocking inflammation can decrease ECM deposition and improve function (*38*). Therefore, we tested next if blocking macrophage expansion with intraperitoneal injections of an antibody to colony-stimulating factor 1 receptor (a-CSF1-R) affects the observed ECM changes in DRGs and functional outcomes in BIPN (Fig. 7A) We focused on males, because the increase in the ECM was male specific, and male mice responded to BTZ with greater immune expansion in DRGs. We first validated that a-CSF1R treatment resulted in depletion of CD68 positive macrophages in DRG compared to the IgG control. (Fig 7B, C). Indeed, while mice treated with BTZ had increased numbers of CD68 positive cells as compared to vehicle treated mice, mice that received a-CSFR1 and BTZ had significantly fewer CD68 positive cells than mice treated with BTZ or vehicle (Fig. 7B, C). As shown earlier (Fig. 6D), mice treated with BTZ and control IgG had increased GFAP immunoreactivity in SGCs as compared to vehicle treated mice, whereas GFAP expression was normalized in mice treated with BTZ and a-CSF1R (Fig. 7D, E). The decreased GFAP expression was associated with decreased collagen Iα deposition in the DRGs from mice treated with BTZ and a-CSFR1 (Fig. 7F, G). Depleting macrophages also prevented BTZ-induced loss of myelinated axons of the sural nerves (Fig. 7H, I) and intra-epidermal nerve fibers (Fig 7J, K). Furthermore, mice treated with a-CSF1R and BTZ as compared to mice treated only with BTZ had significantly improved rotarod performance and nerve conduction studies and trended towards improved mechanical hyperalgesia (Fig. 7L-N). Thus, depleting macrophages resulted in normalization of the DRG microenvironment, prevented axon degeneration and improved functional outcomes in DRGs from male mice treated with BTZ.

**Figure 7:**
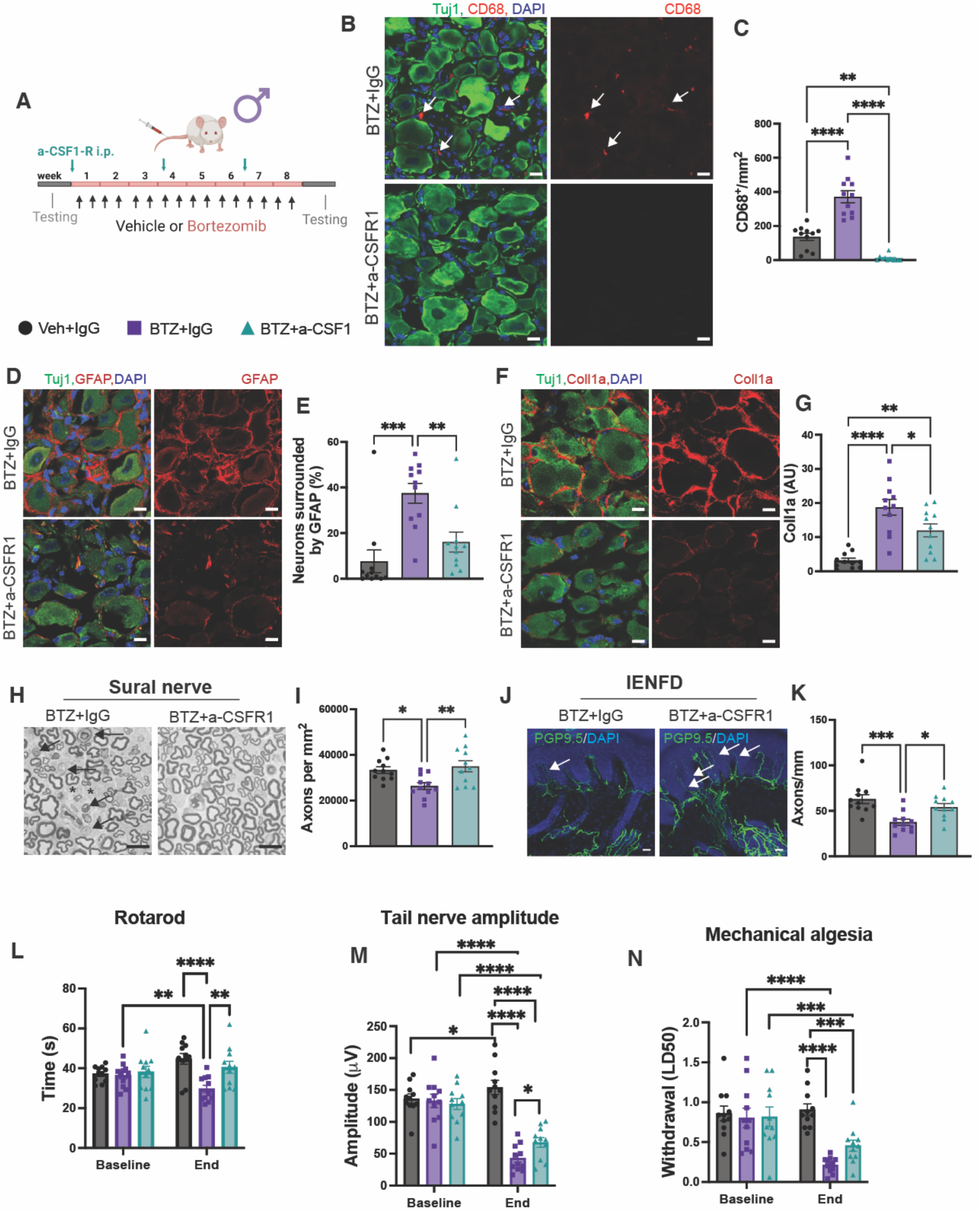
Depleting macrophages prevents bortezomib-induced neuropathy in male mice. **(A)** Experimental layout. **(B)** Photomicrographs of lumbar DRGs stained for tubulin (Tuj1, green, neurons), CD 68 (red, macrophages), and DAPI (blue, nuclei) from mice treated with BTZ (BTZ+ control IgG) or BTZ and a-CSFR1. Note the absence of macrophages after a-CSFR1 treatment. Scale bars 10 μm. **(C)** Quantification of CD68-positive cells in DRGs. One way ANOVA **P=0.0018; ****P<0.0001; n=11 mice per group. **(D)** Photomicrograph of DRGs stained for tubulin (green, neuron), GFAP (red, SGCs), and DAPI (blue, nuclei) from mice treated with BTZ and control IgG or BTZ and a-CSFR1. Scale bars 10 μm. **(E)** Quantification of neurons surrounded more than 50% by GFAP-immunoreactivity; One way ANOVA **P=0.0068; *** P=0.0002. n=11 mice per group (**F**) Photomicrograph of lumbar DRGs stained for tubulin (green, neuron), collagen 1α (coll1a; red), and DAPI (blue, nuclei). Scale bars 10 μm. **(G)** Quantification of collagen-Iα fluorescence intensity in DRGs. One way ANOVA *P=0.0314; **P=0.0040, ****P<0.0001; n=11 mice/group. **(H)** Photomicrographs of toluidine-blue stained semithin sections of cross-sectioned sural nerves from mice treated with BTZ and control IgG or BTZ and a-CSFR1. Note the increased inter-axonal space (asterisks) and degenerating axons (arrows) after BTZ treatment, which was normalized by co-treatment with aCSFR1. Scale bars 10 μm. **(I)** Quantification of axon density of the distal sural nerve. One way ANOVA *P=0.0296, **P=0.0064; n=11 mice per group. **(J)** Photomicrographs of intraepidermal nerve fibers (IENF) of footpads stained for PGP9.5 (green) and DAPI (blue, nuclei). White arrows point to IENF. Scale bars 20 μm. **(K)** Quantification of IENF density; One way ANOVA *P=0.0208, ***P=0.0004, n = average of 3-4 sections per animal, 11 mice/group. (**L, M, N)**: Rotarod (L), compound tail nerve action potential (M), and von Frey testing (N) of mice before (baseline) and immediately after treatment (End). L-N: Two-way ANOVA with Fisher’s LSD; (L) BTZ vs BTZ+aCSFR1 **P=0.0017; Veh baseline vs end **P=0.0036, BTZ baseline vs end **P=0.008; veh vs BTZ ****P<0.0001; (M) BTZ vs BTZ+CSF *P=0.048; Veh baseline -end *P=0.047; ****P<0.0001; (N)***P=0.0009, ****P<0.0001; L-N n=11 mice/group.

## DISCUSSION

Our findings reveal degeneration and dysfunction in both myelinated and unmyelinated peripheral axons in male and female mice in BIPN, with males generally exhibiting a more severe phenotype, which we confirmed in human patients. In the mouse model, BTZ led to significant alterations in the DRG microenvironment, notably marked by extensive transcriptional changes of SGCs characterized by increased expression of genes of the ECM in males. This was accompanied by greater immune cell expansion and increased fibrosis in males compared to females. Systemic administration of anti-CSF1R reduced macrophage numbers in the DRG, normalized GFAP expression in SGCs, normalized the ECM, blocked the degeneration of myelinated and unmyelinated axons and improved functional outcomes, including performance in rotarod tests and nerve conduction studies in male mice.

Sex differences in therapeutic responses to BTZ-induced pain have recently been reported in rodent studies. Antagonists to adenosine receptor 3 and sphingosine 1 phosphate receptor 1 are effective in decreasing BTZ-induced pain in males, but not females, highlighting the importance of sex in the effect of BTZ (*42*). Here, we extend these findings to humans, demonstrating that men had a more severe phenotype than women. BIPN studies in humans rarely consider patient sex, but one study assessing patient-reported outcomes revealed no differences in symptom severity between men and women (*4*), consistent with our finding of no differences in pain intensities between the sexes. However, the results of nerve conduction studies and neurological examination were not reported, and it was these tests that uncovered the important differences detected in the present study. Re-analysis of data from a previously published large cohort of patients with BIPN, in which the total neuropathy score (TNSc) and sex were reported for each individual patient, exposed a significant higher TNSc in males (*9*), indicating more severe BIPN in male than in female patients. Thus, we here uncovered a sex difference in the severity of BIPN both in rodents and humans that has noteworthy implications for research and future drug development, highlighting an urgent need for tailored, sex-specific therapeutic approaches.

Sex differences in peripheral neuropathy are evident in terms of prevalence, severity, symptoms, and treatment responses, with a general female predominance observed for painful peripheral neuropathies (*23–25, 43–45*). Although the underlying mechanisms for these sex disparities are not fully understood, hormonal influences, pain perception, and inflammatory responses are thought to contribute (*46*). Notable variations in DRG gene expression between men and women with neuropathic pain have been identified(*47*), along with sex-specific dimorphisms in DRG nociceptor sensitization (*48*). In this study, we found that SGCs, rather than neurons, exhibited the most pronounced sex-specific gene expression at baseline. Gene ontology analysis reveals an enrichment of pathways related to proteasome function and assembly in males, likely rendering male SGCs more susceptible to the effects of the proteasome inhibitor BTZ. Accordingly, SGCs show the largest transcriptomic response to BTZ, more in males, and resulting in the more severe phenotype of BIPN in males.

SGCs occupy a unique space in the DRG microenvironment. On one side they tightly enwrap each neuron, forming a functional glia-neuron unit that allows for close mutual neuron– SGCs interactions. On the other side, they are in contact with the DRG microenvironment, including macrophages, fibroblasts and endothelial cells. Thus, SGCs are well-positioned to monitor and relay information, and possibly material, from the microenvironment to neurons and vice versa. SGC genes encoding different components of the ECM, including different collagens, ranked among the most upregulated in male mice treated with BTZ. This effect was accompanied by massively increased collagen deposition in the interstitial space. Other DRG cell types also increased expression of genes encoding ECM in response to chronic BTZ, in male, but not female mice. These included fibroblasts, non-myelinating and myelinating Schwann cells, and vascular smooth muscle cells, resulting in a strongly pro-fibrotic state in DRGs of male mice treated with BTZ.

The ECM is a dynamic and interactive three-dimensional network composed of a diverse array of macromolecules (*33*). It not only provides structural support and confers mechanical properties to cells and tissues but also transmits signals to the intracellular environment through receptors such as integrins (*34, 35*). In CNS structures, such as the cortex, hippocampus, and amygdala, the ECM has been shown to limit neuroplasticity by inhibiting activity-dependent structural reorganization of presynaptic and postsynaptic compartments (*34*). Accordingly, disruption of the ECM enhances synaptic plasticity, memory acquisition, and cognitive flexibility. In the hypothalamic arcuate nuclei, increased ECM in the form of perineuronal nets blocks insulin molecules from binding their receptors thereby leading to hyperglycemia and metabolic syndrome, whereas disassembly of the ECM reversed the disease process (*32*). The role of the ECM in the DRG microenvironment lags behind the growing awareness of its the critical importance in overall CNS homeostasis. However, recent discoveries augur an important role for collagen signaling pathways in DRGs(*49*). Collagen signaling pathways ranked highest in analyses of murine DRG scRNAseq data with CellChat, indicating that the ECM contributes the most to predicted interactions occurring in the DRG (*49*). Strikingly, ECM organization in the DRG was the top category for differentially expressed pathways in murine models of inflammatory and neuropathic pain (*50*). In line with these findings, augmented ECM has been observed in DRGs of patients suffering from various pain conditions (*51*). Recent studies indicate that ECM alterations may also be involved in pain induced by the chemotherapeutic paclitaxel(*52*). Paclitaxel treatment results in decreased expression of the tissue inhibitor matrix metalloproteinase 3 in SGCs of male and female mice, which correlates with elevated levels of matrix metalloproteinases 14 and Adam17— enzymes that degrade various ECM components (*52*). While the upregulation of these enzymes suggests a reduction in ECM, we have observed a significant increase in collagen in our model of chronic BIPN. This difference may arise from distinct responses of the ECM to chemotherapeutics with different mechanisms of action (paclitaxel tubulin inhibitor, BTZ proteasome inhibitor) or the shorter duration of paclitaxel treatment (two weeks compared to two months in the present study). Taken together, these findings suggest an important role for collagen pathways in cell-cell communication in DRGs and in the pathophysiology of CIPN.

Fibrosis in peripheral organs is a result of persistent inflammation (*37*). Here, we find an increase of immune cells, including macrophages, in DRGs of mice with BIPN. Through their potential pro-fibrotic and pro-inflammatory properties, macrophages are at the crossroad of key pathogenic processes and associated manifestations. We identified macrophages in DRGs with both pro-inflammatory and pro-fibrotic gene expression patterns. After macrophages were depleted by blocking the CSF1-receptor, we observed less SGC activation, and decreased collagen deposition in the DRGs of BTZ-treated mice. This normalization of the DRG microenvironment was associated with preservation of axons and improved functional outcomes.

Our inability to restrict macrophage depletion to the DRG is a limitation of this study. Selective *in vivo* depletion of a subpopulation defined by multiple markers within identified, intended CNS or PNS structures remains challenging at this time, rendering such proof elusive. Therefore, we cannot rule out that depletion of macrophages in other parts of the nervous system, such as, e.g., the peripheral nerve, contributes to the prevention of BIPN. Nevertheless, we identified a novel mechanism of BIPN and a target to prevent it. Depletion of macrophages with aCSFR1 has been shown to decrease multiple myeloma burden by itself (*53, 54*), an effect that was increased with the addition of BTZ (*54*) suggesting that targeting macrophages is a promising avenue to both increase cancer targeting efficacy of BTZ and at the same time reduce the occurrence of BIPN. Insofar as global depletion of macrophages is unlikely to be translatable to the clinic, a more practical goal would be to identify and target molecules that activate macrophage subpopulations and other effectors that we know lead to neuropathy. The work described here provides the necessary foundation that will permit pursuit of this kind of approach.

To summarize, we have discovered that chronic BTZ causes inflammation, SGC reactivity and neurofibrosis in DRGs, more prominently in males than females. Blocking inflammation significantly reverses the effect. Targeting the DRG microenvironment thus is a novel and promising strategy to treat BIPN and possibly other neuropathies

## MATERIALS AND METHODS

### Study design

The primary objective of this research was to identify mechanisms of BIPN. We evaluated this hypothesis by creating a mouse model of BIPN that replicates route of administration and long duration of BTZ in human patients. To keep the groups balanced, animals were selected into the vehicle group or the BTZ treatment group based on pre-treatment results. All animal studies were performed by investigators blinded to treatment groups. Power analyses based on pilot and published data from our laboratory indicated that *n* = 10 to 15 mice per group would be needed to yield 80% power with α set to 0.05. Four cohorts of either female or male mice (each cohort consisting of 10 to 15 mice per group) were used to generate the datasets. Data were analyzed for each cohort separately and then combined. If different results were obtained between the cohorts, the data are reported for each cohort. To evaluate sex differences in BIPN in humans, we evaluated data from patients enrolled into the Peripheral Neuropathy Research Registry and from the Neuromuscular clinic at Washington University St. Louis with IRB approval.

### Animals

All procedures were approved by the Washington University School of Medicine in Saint Louis Institutional Animal Care and Use Committee (Protocols #20180196 and #210348). This manuscript was prepared in adherence to the ARRIVE guidelines.

Mice were housed and cared for in the Washington University School of Medicine animal care facility, which is accredited by the Association for Assessment & Accreditation of Laboratory Animal Care (AALAC) and conforms to the PHS guidelines for Animal Care. Male and female Balb/cJ mice (Jackson), ranging from 8 to 12 weeks of age at the beginning of the experiment were treated twice weekly either with vehicle (5% EtOH, 5% Tween-80, 90% sterile saline) or BTZ dissolved in vehicle at a dose of 0.8 mg/kg (Millipore Sigma) by tail vein injection for 8 weeks. Mouse monoclonal antibody to colony stimulating factor 1 receptor (CSFR1; BioXCell) or mouse monoclonal rat IgG2a isotype control (BioXCell) were administered at 1.5 mg intraperitoneally every three weeks. All animals underwent nerve conduction studies and behavioral testing during the two weeks before and after the last injection as previously described (*55*; supplementary methods).

### Tissue Preparation

Mice were deeply anesthetized and euthanized either by controlled CO2 inhalation or by transcardial perfusion of 4% PFA in PBS.

*Toluidine blue staining and axon quantification* The sural nerves and L4/5 DRGs were fixed by immersion in 3% glutaraldehyde and processed as previously described (*55*). Briefly, after incubation in osmium tetroxide (Sigma), tissue was dehydrated and embedded in araldite/DDSA/DMP30 (Electron Microscopy Sciences). Tissue was sectioned (400 nm thick) on a Leica UC7 ultramicrotome and stained with 1% toluidine blue (Fisher Scientific). Sections were imaged using a 63x oil immersion on a Zeiss LSM 800 microscope. Photos were stitched in Zeiss Zen Blue software and axon number and density of distal sural nerves was analyzed by counting all normal appearing and degenerating axons in cross sections using ImageJ. The area of the cross section was measured in ImageJ and the total number of axons per mm^2^ calculated.

*Immunohistochemistry* Fixed, free floating 50-µm thick sections of footpads from the hind limbs were stained with rabbit anti-Protein Gene Product 9.5 (1:500, Lifespan Biosciences), followed by Cy3- or AF488 conjugated secondary antibody (Invitrogen, 1:500). Sections were mounted and coverslipped using Prolong Gold with DAPI (Molecular Probes). PGP 9.5 positive intraepidermal nerve fibers (IENFs) crossing into the epidermis were counted using 40x objective on a ZEISS LSM 800 microscope. IENF densities were averaged from three to four sections for each animal. Counting was performed with the observer blinded to the treatment group. Mounted sections of frozen DRGs were stained with in rabbit anti-GFAP (1:500 Dako), chicken anti-tubulin (1:1000 Abcam), rabbit anti-Fabp7 (1:1,000, ThermoFisher), rabbit anti-KCNJ10 (1:1,000 Alomone), rat anti-CD68 (1:500, Bio Rad), which was followed by secondary antibodies Cy3 or AF488 conjugated goat anti-rabbit, goat anti-rat, or goat anti-chicken (all 1:500, Invitrogen). Slides were coverslipped in ProlongGold with DAPI (Molecular Probes). DRG were imaged and total neurons, neurons surrounded by GFAP, or CD68 positive cells were counted and the area of the DRG determined in ImageJ.

### Single Cell RNAseq

*Sample collection and library prepration* was done as previously described (*31*). Briefly, L4 and L5 DRG’s were collected, ganglia mechanically dissociated to a single-cell suspension and cells were FACS sorted using MoFlo HTS with Cyclone (Beckman Coulter, Indianapolis, IN). Single-cell RNA-Seq libraries were prepared using GemCode Single-Cell 3′ Gel Bead and Library Kit (v3, 10 x Genomics) and sequenced on an Illumina NovaSeq 6000 platform.

*Read alignment and processing* Raw sequencing data in FASTQ format were processed using Cell Ranger v7.0 (10x Genomics) to generate digital gene expression matrices for each sample.

*Integration and clustering* Unfiltered expression matrices generated by Cell Ranger were analyzed using the Seurat v5.0 R package (*56*). Cells with fewer than 500 genes detected were excluded from analysis. Data normalization was performed using the SCTransform method (*57*). Principal component analysis (PCA) was conducted using the top 50 principal components and integration of seven original batches was performed using the reciprocal PCA (RPCA) method implemented in Seurat. Clustering was performed using a shared nearest neighbor (SNN) modularity optimization-based algorithm with a resolution parameter of 0.7, which was determined after examining multiple resolutions with the Clustree R package (*58*). Cluster identity was assigned based on the expression of major cell type marker genes. Low-quality non-neuronal clusters were removed if they met the following criterion: average number of genes detected (nFeature_RNA) < 1400. Clusters with enriched expression of neuronal markers were subset and clustered separately from non-neuronal cells to identify subtype. Low-quality neuron clusters were removed if they met any of the following criteria: significant co-expression of non-neuronal cell type markers, average nFeature_RNA < 1000, or fewer than 10 cells total in at least one of the four treatment groups. Cluster identities were assigned based on published marker gene enrichment (*27*) and transferred to the main Seurat object, with low-quality cells removed.

*Marker gene and differential gene expression algorithms/statistics* Differentially expressed genes (DEGs) and cluster marker genes were identified using the Seurat FindMarkers function, which employs a Wilcoxon rank sum test with bonferroni correction.

*Subclustering* Immune cells and mitotic cells were subset from the main object and subjected to reclustering following the same procedure as described for the main object to identify subpopulations within each group.

*Gene ontology* Gene Ontology (GO) enrichment analysis was performed using the clusterProfiler R package (*59*).

*CellChat* Cell-cell communication analysis was conducted using CellChat v2.0 (*36*). The analysis compared bortezomib-treated (BTZ) and vehicle-treated (VEH) conditions, with male and female data combined for each condition. CellChat was run with default parameters to identify changes in intercellular communication networks between treatment groups. The algorithm infers cell-cell communication by analyzing the expression levels of ligand-receptor pairs and their cofactors across different cell populations, using a probability-based approach.

### Human data

Data of patients enrolled into the Peripheral Neuropathy Research Registry and from the neuromuscular clinic at Washington University St. Louis were evaluated. The Peripheral Neuropathy Research Registry (PNRR) is an ongoing multi-center collaboration seeking to characterize the phenotypes of acquired peripheral neuropathies in defined patient cohorts (*22*). The Institutional Review Board (IRB) at each consortium member has approved the PNRR protocol and all enrolled patients provided informed consent. The diagnosis of BIPN was based on typical symptoms (including neuropathic pain, numbness, muscular weakness, balance impairment, and autonomic symptoms), neurological examination by a board-certified neuromuscular specialist, nerve conduction studies and/or skin biopsy. Onset of symptoms had to correlate with the timing of receiving BTZ. Patients completed a questionnaire regarding their neuropathic symptoms during the past 7 days. Symptoms commonly associated with peripheral neuropathy like pain, numbness, muscular weakness, balance impairment, autonomic symptoms, sleep difficulties, and muscle cramping were captured. To increase group sizes, additional patients were identified from the Neuromuscular Center at Washington University St. Louis, and clinical information was obtained by reviewing medical records. All procedures were approved by the human studies and ethics committee at Washington University in St. Louis.

### Statistical Analysis

Data collection and analyses were performed blind to the conditions of the experiments. Single-cell RNAseq analysis was performed in an unbiased manner using established algorithms. Unless stated otherwise, the data are reported as means ± the standard error of the mean (SEM). Comparisons between groups was made with one-way and two-way ANOVA, unpaired *t*-test, or Chi-square tests as warranted. Two-sided significance tests were applied when appropriate and a *P* ˂ 0.05 was considered statistically significant. Statistics were calculated using GraphPad Prism10.

## List of Supplementary Materials

Materials and Methods

Fig. S1 to S4

Table S1

Data file S1

Data file S2

## Funding

National Institutes of Health grant R37CA267905 (SG), R35 NS122260 (VC), R01 NS111719 (VC)

Pilot Project Award from the Hope Center for Neurological Disorders at Washington University (SG, VC)

Siteman Cancer Center (SG)

Sims Family Fund for Neurological Research (SG)

## Author contributions

Conceptualization: MBT, VC, SG

Methodology: MBT, AS, CNT, OA, SXY, SG

Software: MBT

Investigation: MBT, AS, CNT, VDG, NFS, OA, SXY, SK, SSG, SL, AP, SG

Formal analysis: MBT, AS, CNT, NFS, OA, SK, SSG, SG

Visualization: MBT, AS, CNT, NFS, SL, SG

Data curation: MBT

Funding acquisition: VC, SG

Project administration: SG Supervision: VC, SG

Writing – original draft: SG

Writing – review & editing: MBT, AS, CNT, VDG, NFS, OA, SXY, SK, SSG, SL, AP VC, SG

## Data and materials availability

Raw and processed scRNA sequencing data have been deposited in the Gene Expression Omnibus (accession number GSE275255). Additional data presented in the main text or the Supplementary Materials are available upon reasonable request from the corresponding author.

