## Supplementary Materials for "Macrophage depletion restores the DRG microenvironment and prevents axon degeneration in bortezomib-induced neuropathy"

**Supplementary Methods**

*Nerve conduction studies*: Recording electrodes were placed subcutaneously at the base of the tail and stimulating electrodes were inserted distally 30 mm from the negative recording electrode. Evoked compound nerve action potentials (CNAPs) were acquired using a Viking Quest electromyography machine (Nicolet). Responses to 15 stimuli were averaged. The amplitude was measured from baseline to first peak and conduction velocity calculated as a function of distal latency and distance between stimulating and recording electrode (30 mm).

*Von Frey*: Animals were habituated in clear Plexiglass boxes on a raised wire mesh for 4 h. Once mice were resting and not exploring, testing was performed according to the up-down method *(60)*. The withdrawal threshold was calculated for each paw and averaged between the two *(61)*. Von Frey experiments prior to and following treatment were performed at the same time of day by the same experimenter blind to the condition.

*Rotarod* Rotarod experiments were conducted as previously described *(55)*. Briefly, after acclimatizing to the testing room, animals were placed on the rotarod (Columbus Instruments Rotamex 5) at a constant speed of five rotations per minute (rpm) for 2 min for a total of 5 trials with 10 minutes of rest between trials. Then, animals were placed on the rotarod and the rotation speed was increased by 3 rpm every 10 s. The time to fall off the rod was recorded over five trials and averaged.

*Tissue processing*: After incubation overnight in 1% osmium tetroxide (Sigma), tissue was dehydrated into 100% propylene oxide. Nerves were then placed into ascending concentrations of araldite/propylene oxide solutions and embedded in araldite/DDSA/DMP30 (Electron Microscopy Sciences) and polymerized overnight at 60 °C. Nerves were cross sectioned (400 nm thick) on a Leica UC7 ultramicrotome and stained with 1% toluidine blue/2% sodium borate (Fisher Scientific). Sections were imaged using a 63x oil immersion on a Zeiss LSM 800 microscope. Photos were stitched in Zeiss Zen Blue software and axon number and density of distal sural nerves was analyzed by counting all normal and degenerating axons in the entire cross section using ImageJ. The area of the cross section was measured in ImageJ and the total number of axons per $mm^2$ calculated. DRGs were similarly cross sectioned and stained with 1% toluidine blue/2% sodium borate and imaged using 63x oil immersion.

*Immunohistochemistry and determination of intraepidermal nerve fiber density* The plantar surface of each foot was fixed in Zamboni's fixative (0.12% picric acid, 2% paraformaldehyde in 0.1 M phosphate buffered saline (PBS) over night, cryoprotected and frozen. Four contiguous series of 50-µm thick cross sections were cut on the cryostat (Leica CM1950) and collected such that the entire plantar surface was sampled at 200 µm intervals. Free floating sections were stained with rabbit anti-Protein Gene Product 9.5 (1:500, Lifespan Biosciences) overnight, followed by Cy3-conjugated or AF488 conjugated anti-rabbit secondary antibody (Invitrogen; 1:500) for 2 h. Sections were mounted and coverslipped using ProlongGold with DAPI (Molecular Probes) to allow visualization of nuclei. PGP 9.5 positive intraepidermal nerve fibers (IENFs) crossing into the epidermis were counted using 40x objective on a ZEISS LSM 800 microscope. IENF densities were averaged from three separate sections for each animal. Counting was performed with the observer blinded to the treatment group.

*Immunohistochemistry of DRGs* After perfusion with 4% PFA, lumbar DRGs were dissected out and placed in 4% PFA over night before cryoprotected and frozen. Eight-µm thick, mounted sections were stained with in rabbit anti-GFAP (1:500 Dako), chicken anti-tubulin (1:1000, Abcam), rabbit anti-Fabp7 (1:1000, ThermoFisher), rabbit anti-KCNJ10 (1:1000, Alomone labs), or rat anti-CD68 (1:250, BioRad). Secondary antibodies were Cy3- or AF488 conjugated goat anti-rabbit, or goat anti-chicken, goat anti-rat (all 1:500; Invitrogen). Slides were coverslipped in ProlongGold with DAPI (Molecular Probes).

DRGs were imaged at 20x or 40x on a Zeiss LSM800, and images were stitched using Zeiss Zen Blue software. Total neurons, neurons surrounded by GFAP-immunoreactivity or CD68 positive cells were counted and the area of the DRG determined in ImageJ. The averages from three separate sections of each DRG were calculated.

**Single Cell RNAseq**

*Sample collection and library prepration* L4 and L5 DRG's were collected into cold Hank's balanced salt solution (HBSS) with 5 % Hepes, then transferred to warm Papain solution and incubated for 20 min in 37 °C. DRG's were washed in HBSS and incubated with Collagenase for 20 min in 37 °C. Ganglia were then mechanically dissociated to a single-cell suspension by triturating in culture medium (Neurobasal medium), with Glutamax, PenStrep and B-27. Cells were washed in HBSS+ Hepes + 0.1% BSA solution, passed through a 70 micron cell strainer. Hoechst dye was added to distinguish live cells from debris and cells were FACS sorted using MoFlo HTS with Cyclone (Beckman Coulter, Indianapolis, IN). Sorted cells were washed in HBSS+ Hepes + 0.1% BSA solution and manually counted using a hemocytometer. Solution was adjusted to a concentration of 500 cell/microliter and loaded on the10 X Chromium system. Single-cell RNA-Seq libraries were prepared using GemCode Single-Cell 3′ Gel Bead and Library Kit (v3, 10 x Genomics) and sequenced on an Illumina NovaSeq 6000 platform.

#### Supplementary figures

#### Figure S1

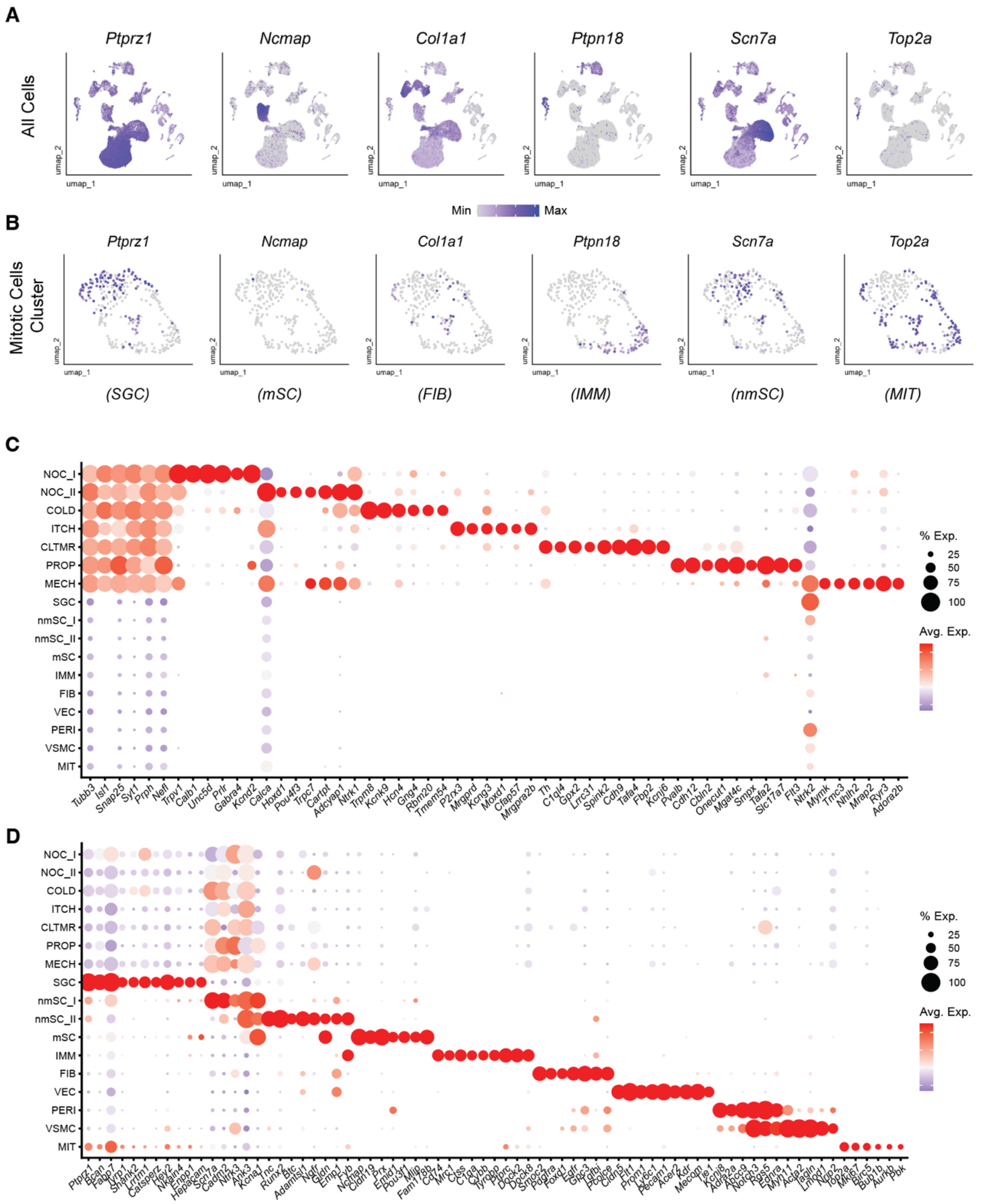


**Supplementary figure 1**: **Marker gene expressions of identified cell types in the DRG**. **(A, B)** Marker gene expression UMAP plots in all major cell types (A) or mitotic cells only (B). Bottom: cell type associated with each marker gene. **(C)** Dotplot of neuron cluster-specific marker genes **(D)** Dotplot of non-neuronal cluster-specific marker genes.

### Figure S2

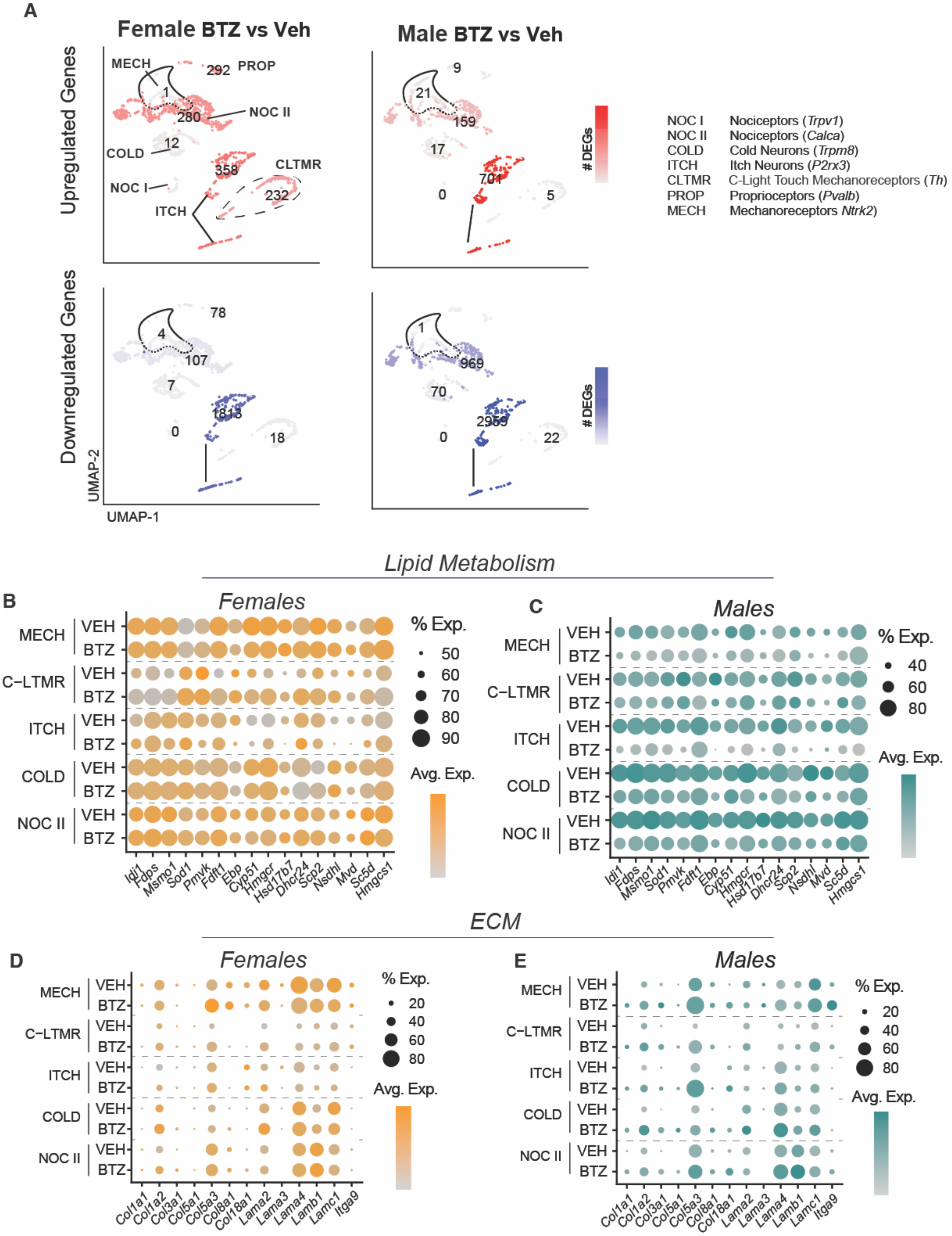

A
Female BTZ vs Veh
Male BTZ vs Veh
Upregulated Genes
Downregulated Genes
MECH
PROP
NOC II
COLD
CLTMR
NOC I
ITCH
UMAP-2
UMAP-1
### DEGs
NOC I Nociceptors (Trpv1)
NOC II Nociceptors (Calca)
COLD Cold Neurons (Trpm8)
ITCH Itch Neurons (P2rx3)
CLTMR C-Light Touch Mechanoreceptors (Th)
PROP Proprioceptors (Pvalb)
MECH Mechanoreceptors Ntrk2)
Lipid Metabolism
B
Females
C
Males
MECH
C−LTMR
ITCH
COLD
NOC II
VEH
BTZ
% Exp.
Avg. Exp.
Idi1
Fdps
Msmo1
Sod1
Pmvk
Fdft1
Ebp
Cyp51
Hmgcr
Hsd17b7
Dhcr24
Scp2
Nsdhl
Mvd
Sc5d
Hmgcs1
ECM
D
Females
E
Males
Col1a1
Col1a2
Col3a1
Col5a1
Col5a3
Col8a1
Col18a1
Lama2
Lama3
Lama4
Lamb1
Lamc1
Itga9

**Supplementary figure 2: Differential gene expression in DRG neurons of female and male mice with BIPN**. **(A)** UMAP plots displaying differential gene expression magnitude in the neuronal cell type clusters from female and male samples depicting upregulated (Red, upper) and downregulated (blue, lower) genes (p-adj < 0.05). Numbers indicate differentially expressed genes in each cell cluster. **(B-E)** Gene Ontology enrichment analysis of differentially expressed lipid metabolism or (B, C) or ECM (D, E) genes (p-adj < 0.05, log2FC > 0.25) in different neuron population from female (B, D) and male (C, E) mice treated with vehicle or BTZ.

**Figure S3**

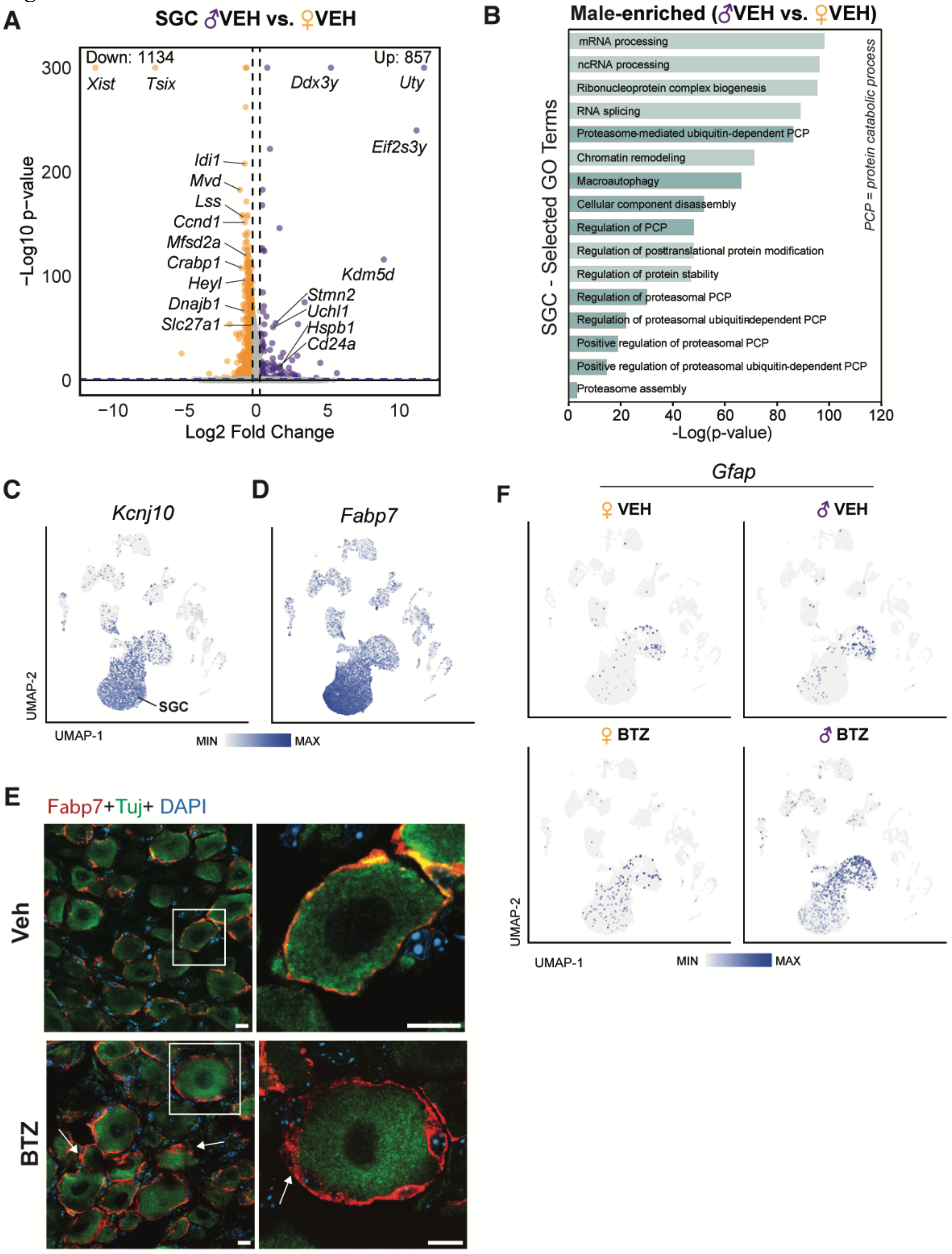
A
SGC ♂VEH vs. ♀VEH
Down: 1134
Up: 857
Xist
Tsix
Ddx3y
Uty
Eif2s3y
Idi1
Mvd
Lss
Ccnd1
Mfsd2a
Crabp1
Heyl
Dnajb1
Slc27a1
Kdm5d
Stmn2
Uchl1
Hspb1
Cd24a
−Log10 p−value
Log2 Fold Change
B
Male-enriched (♂VEH vs. ♀VEH)
mRNA processing
ncRNA processing
Ribonucleoprotein complex biogenesis
RNA splicing
Proteasome-mediated ubiquitin-dependent PCP
Chromatin remodeling
Macroautophagy
Cellular component disassembly
Regulation of PCP
Regulation of posttranslational protein modification
Regulation of protein stability
Regulation of proteasomal PCP
Regulation of proteasomal ubiquitin-dependent PCP
Positive regulation of proteasomal PCP
Positive regulation of proteasomal ubiquitin-dependent PCP
Proteasome assembly
SGC - Selected GO Terms
PCP = protein catabolic process
-Log(p-value)
C
Kcnj10
SGC
UMAP-2
UMAP-1
D
Fabp7
MIN
MAX
E
Fabp7+Tuj+ DAPI
Veh
BTZ
F
Gfap
♀ VEH
♂ VEH
♀ BTZ
♂ BTZ
UMAP-2
UMAP-1
MIN
MAX

**Supplementary figure 3: Differential gene expression in satellite glial cells of female and male mice treated with vehicle. (A)** Volcano plot of differentially expressed genes of SGCs from vehicle treated male and female mice yellow: female enriched. Purple: Male enriched. **(B)** Gene Ontology enrichment analysis of differentially expressed genes (p-adj $< 0.05$, log2FC $> 0.25$) in SGCs from males versus females treated with vehicle shows enrichment in proteasome processing in males (dark teal bars). **(C, D)** UMAP plots demonstrating gene expression of Kcnj10 (C) and Fabp7 (D) predominantly in SGCs. **(E)** Photomicrographs depicting SGCs from males treated with vehicle or BTZ stained with tubulin (Tuj, green, neurons) Fabp7 (red, SGCs) and DAPI (blue, nuclei). Note widening of the Fabp7-positive perineuronal cell-layer after BTZ (white arrows). Right column higher magnification of boxed area in photos to the left. Sclae bars 10 µm. **(F)** UMAPS of GFAP gene expression demonstrating increased GFAP in SGCs from female and male mice with BIPN.

#### Figure S4

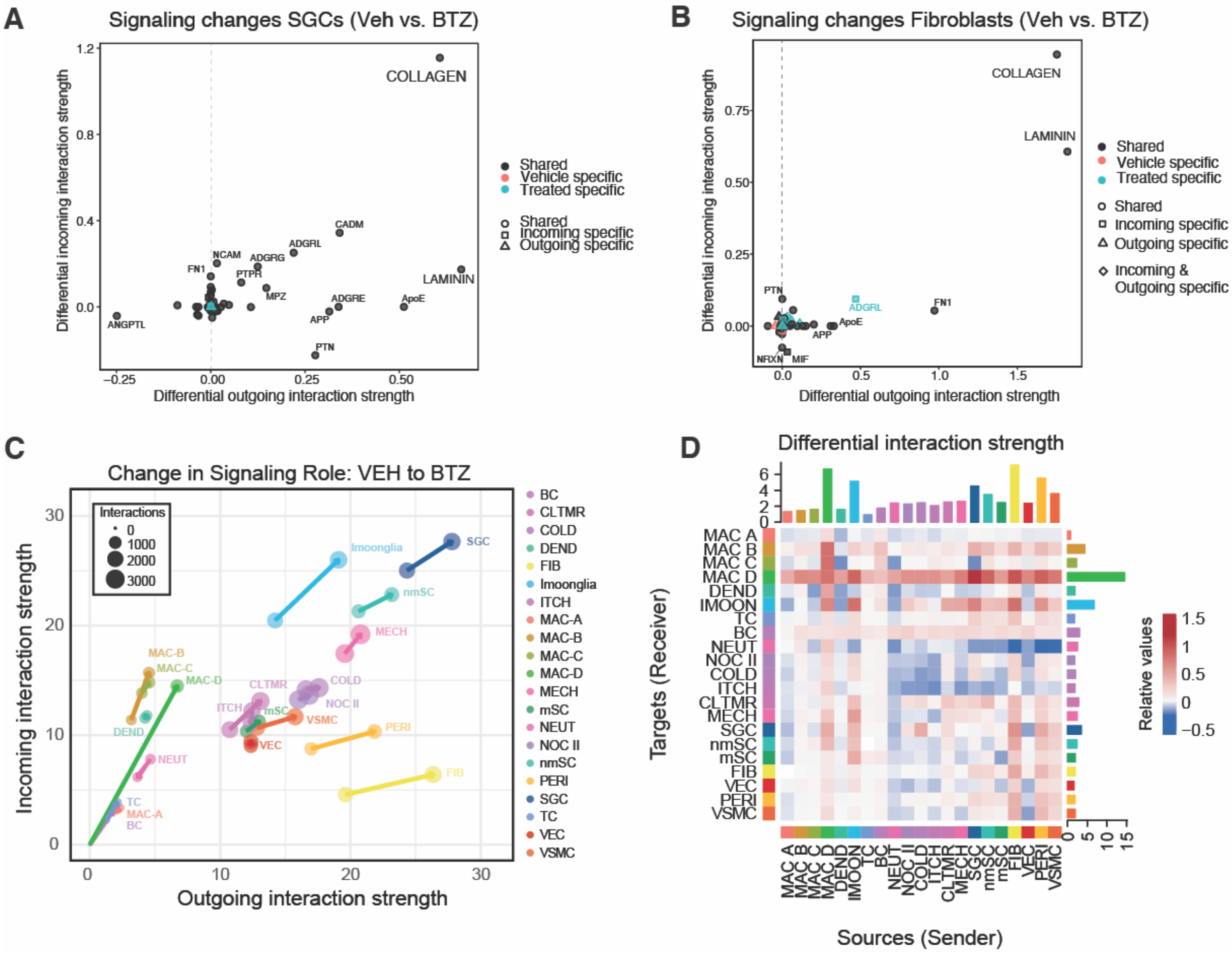


**Supplementary figure 4**: **Signaling changes in DRGs in BIPN**. (**A, B**): Cell chat analysis demonstrating that both incoming and outgoing communication through the COLLAGEN and LAMININ pathways increases in SGCs (A) and fibroblasts (B) following BTZ. These pathways are also present in controls (grey color). (**C**): The signaling strength of several cell types in the DRG increases following BTZ. Arrow points to direction of change (increase up, decrease down). Y axis- change in incoming interaction strength, X axis change in outgoing interaction strength. Macrophage D profoundly increases incoming but also outgoing interaction strengths, whereas fibroblasts show especially increase of outgoing interaction strength in BIPN. SGC is the cell type in DRG with the highest interaction strength at baseline, which increases further (both incoming and outcoming) following BTZ. **(D):** Differential interaction strength between individual cell types with receivers plotted on the y axis and source on the x axis.

**Supplementary Table**

| | Male (n = 24) | Female (n = 16) | P-Value |
|---|---|---|---|
| **Age, median (min, max) *** | 67.5 (41,80) | 67.5 (47, 83) | 0.7773 |
| **Race, n (%) **** | | | |
| *White* | 18 (75.0%) | 12 (75.0%) | >0.9999 |
| *Non-White* | 6 (25.0%) | 4 (25.0%) | |
| **Comorbidities, n (%) **** | | | |
| *Diabetes Mellitus* | 4 (16.7%) | 2 (12.5%) | >0.9999 |
| *Hyperlipidemia* | 11 (45.8%) | 6 (37.5%) | 0.7471 |
| *Thyroid Disease* | 3 (12.5%) | 3 (18.8%) | 0.6678 |
| *Fibromyalgia* | 1 (4.2%) | 2 (12.5%) | 0.553 |
| *Vitamin B12 Deficiency* | 3 (12.5%) | 0 (0%) | 0.2615 |
| *Autoimmune* | 2 (8.3%) | 2 (12.5%) | >0.9999 |
| *Kidney Disease* | 2 (8.3%) | 5 (31.3%) | 0.0942 |
| *Irritable Bowel Disease* | 1 (4.2%) | 1 (6.25%) | >0.9999 |

*The parametric p-value is calculated by the T-test for numerical variables.
** The non-parametric p-value is calculated by the Fisher's exact test for categorical variables.
